# Morphological profiling reveals neuroprotection via mitochondrial uncoupling in human dopaminergic neurons

**DOI:** 10.1101/2024.09.19.613945

**Authors:** Vyron Gorgogietas, Amélie Weiss, Loïc Cousin, David Hoffmann, Karen Schmitt, Arnaud Ogier, Peter A. Barbuti, Bruno F.R. Santos, Ibrahim Boussaad, Annika Wittich, Andrea Zaliani, Ole Pless, Rejko Krüger, Peter Sommer, Johannes H. Wilbertz

**Affiliations:** Luxembourg Centre for Systems Biomedicine (LCSB), Esch-sur-Alzette, Luxembourg; Ksilink, Strasbourg, France; Department of Neurology, Columbia University Irving Medical Center, New York, NY 10032, USA; Disease Modelling and Screening Platform, Esch-sur-Alzette, Luxembourg; Fraunhofer Institute for Translational Medicine and Pharmacology ITMP, ScreeningPort, Hamburg, Germany; Centre Hospitalier de Luxembourg (CHL), Luxembourg, Luxembourg; Transversal Translational Medicine, Luxembourg Institute of Health (LIH), Strassen, Luxembourg; Parkinson’s Research Clinic, Centre Hospitalier de Luxembourg (CHL), Luxembourg, Luxembourg

## Abstract

Parkinson’s disease (PD) involves multiple pathological processes in midbrain dopaminergic (mDA) neurons, including protein degradation defects, vesicular trafficking disruption, endolysosomal dysfunction, mitochondrial issues, and oxidative stress. Current PD models often lack complexity and focus on single phenotypes. We used patient-derived SNCA triplication (SNCA-4x) and isogenic control (SNCA-corr) mDA neurons, applying high-content imaging-based morphological profiling to identify and rescue multiple phenotypes. Screening 1,020 compounds, we identified top-scoring compounds that restored healthy profiles in SNCA-4x neurons, increasing Tyrosine hydroxylase (TH) and decreasing α-synuclein (αSyn) levels. Several hits were linked to mitochondrial biology. Tyrphostin A9, a mitochondrial uncoupler, and several of its structural analogues decreased ROS levels, normalized mitochondrial membrane potential, and increased respiration. Western blotting confirmed that Tyrphostin A9 reduces αSyn levels. Our study highlights the neuroprotective potential of mild mitochondrial uncoupling in mDA neurons.

## Introduction

Parkinson’s disease (PD), with its diverse phenotypic, neuropathological, and genotypic manifestations, is increasingly acknowledged as a complex disorder rather than a singular disease entity ^1–3^. While traditional models have been instrumental in devising symptomatic therapies to address PD’s motor symptoms, they have been less successful in developing neuroprotective strategies ^4^. Genetic stratification of patients into subgroups has emerged as a crucial approach ^5,6^. Over the past 25 years, significant progress has been made in identifying genes associated with PD, including those responsible for rare monogenic forms of the disease ^2^. These monogenic forms have proven invaluable for PD research, as patient-based cell models exhibit disease-specific cellular phenotypes that mirror those found in postmortem brain tissue ^7–11^. This approach suggests that targets for pharmaceutical interventions might be identifiable based on rare but potent molecular signatures ^12–14^.

Point mutations, as well as duplications and triplications of the wild-type alpha-synuclein gene (*SNCA*), are responsible for causing autosomal dominantly inherited PD ^15–18^. Patients with *SNCA* triplication (SNCA-4x) express α-synuclein protein (αSyn) from four alleles resulting in a roughly 2-fold increase of αSyn, and typically exhibit an early disease onset, accelerated disease progression, and significant dementia. In contrast, patients with SNCA duplications often display a more conventional late-onset PD phenotype. The severity of clinical trajectories in SNCA multiplication appears to correlate with gene dosage. Patient-derived induced pluripotent stem cells (iPSC) carrying the *SNCA* triplication have been successfully differentiated into midbrain dopaminergic (mDA) neurons and organoids ^19–22^. Despite their young age, iPSC mDA neurons exhibit PD-relevant phenotypes. *SNCA* gene triplication leads to an increase in αSyn levels resulting in oligomeric αSyn pathology and elevated αSyn extracellular release ^19,21,23,24^. Multiple pathophysiological defects correlate with increased αSyn protein expression, such as altered lysosomal and mitochondrial functions, dysregulated autophagy increased endoplasmic reticulum stress, and heightened oxidative stress levels ^20,22,24–28^. Due to the interconnection of many of the identified phenotypes, it is currently an open question whether a rescue of one key or multiple cellular PD hallmarks is necessary to achieve a significant neuroprotective effect. Imaging-based profiling is a strategy that transforms the rich data in biological images into multidimensional profiles of extracted features, which can be analyzed to reveal biological signatures, aiding in disease understanding and drug discovery, with advancements in machine learning and computational technologies enhancing its potential ^29^. By capturing the observable characteristics of cells under different conditions, morphological profiling can provide valuable insights into the complex cellular mechanisms underlying PD ^8,9,11^.

In this study, we find that *SNCA* gene triplication results in a multidimensional alteration of the morphological profile in mDA neurons. We further identify multiple small molecular compounds correcting this altered profile to align with the phenotype of isogenic controls (SNCA-corr) mDA neurons. One of the identified compounds, Tyrphostin A9, exerts its neuroprotective effect via mitochondrial uncoupling, which in turn reduces reactive oxygen species (ROS) and high molecular weight αSyn species. In summary, we show that screening small molecules in a PD-relevant human model system using a multi-phenotypical readout can identify modulators of not just a single, but multiple disease relevant phenotypes.

## Results

### SNCA-4x mDA neurons have a distinct but modulable morphological profile

We differentiated patient-derived SNCA gene triplication and isogenic control iPSCs into mDA neurons (SNCA-4x and SNCA-corr) for 30-37 days. These neurons showed the expected mDA neuron morphology and no *de novo* copy number variations. Both cell lines expressed neuro-typical markers β-tubulin III (90-94%), MAP2 (87-89%), and the dopaminergic neuron markers Tyrosine hydroxylase (TH) (84-88%) and FOXA2 (84-90% in TH positive cells) (**Figure S1A-C**). Western blotting revealed increased αSyn protein level and signs of αSyn aggregation in SNCA-4x mDA neurons after D70 (**Figure S1D**). A JC-1 dye assay indicated reduced mitochondrial membrane potential in D35 SNCA-4x mDA neurons (**Figure S1E**). For morphological profiling, mDA neurons were cultured until D36 in 384-well plates and stained with Hoechst, and antibodies against TH, αSyn, and MAP2 (**Figure 1A**). Automated segmentation and feature quantification were used to overcome challenges in assessing subtle phenotypes. Image segmentation was performed on 16 fields per well and 127 morphological features were determined and expressed as means per well. Features were calculated for the whole cell, nucleus, cytoplasm or membrane and describe different aspects of the images (**Figure 1B**, **Table S4**). Texture features capture the spatial distribution and arrangement of pixel intensities in an image. Context features provide information about the spatial relationship between signals, for example the correlation between image channels. Intensity features measure the fluorescence intensity and can provide insights into the expression levels of proteins. Count and area/shape features quantify the number, size and morphology of the cellular components. First, we examined basic morphological features of SNCA-4x and SNCA-corr mDA neurons. SNCA-4x neurons had fewer live cells and approximately 50% increased αSyn protein expression. In our study, we utilized Prostratin as a proof-of-concept molecule due to its proposed activity in degrading αSyn. Previous research has demonstrated that specific phorbol esters, such as PEP005 or Prostratin, can reduce αSyn through proteasomal degradation ^7^. Based on these findings, we hypothesized that Prostratin’s activity would lead to an altered morphological profile in our experimental model. Treatment with Prostratin indeed reduced dead cells and αSyn expression in SNCA-4x neurons to levels similar to SNCA-corr neurons (**Figure 1C**). Neuronal morphological profiles before and after compound treatment were created by scaling feature values and calculating mean values across wells (**Figure 1D**). We used the mean absolute difference to find the features that differed the most between profiles. Comparing SNCA-corr and SNCA-4x neurons, the radial αSyn intensity profile, which measures fluorescence intensity as a function of distance from a central point, differed most. Prostratin-treated and untreated SNCA-4x neurons also varied in number of nuclei, total MAP2 cellular surface, and αSyn staining contrast (**Figure 1E**). These findings show that small molecular treatment can rescue morphological features in mDA neurons which can be measured by morphological profiling.

**Figure 1:**
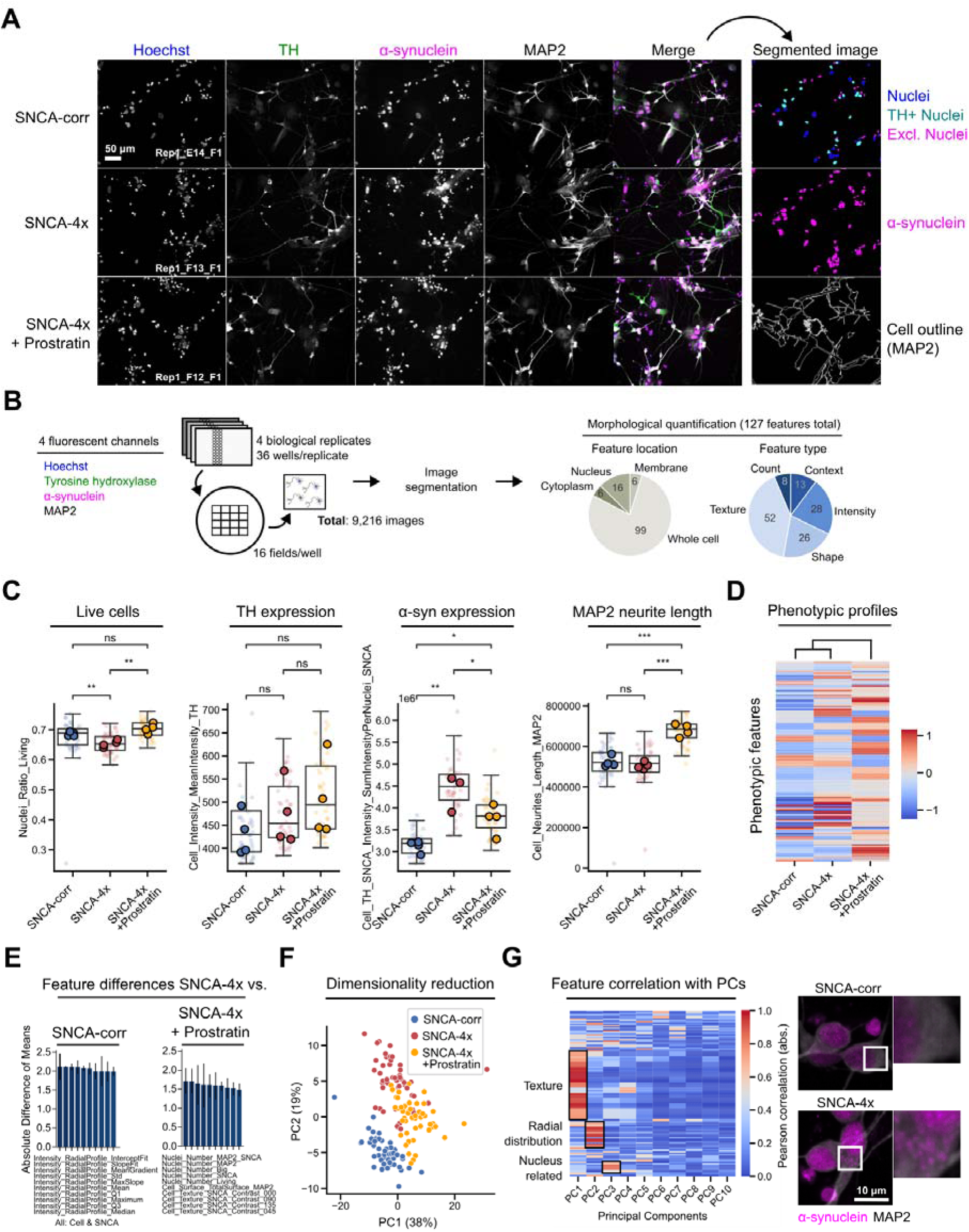
SNCA-4x mDA neurons have a distinct but modulable morphological profile. (A) SNCA-4x and SNCA-corr mDA neurons were differentiated until D33, treated with 5 μM Prostratin or DMSO for 72 hours and immunofluorescently stained. Image channels were segmented for morphological feature quantification. (B) Schematic overview of image data generation and morphological feature types. (C) Raw morphological feature values in control, and untreated and Prostratin treated SNCA-4x mDA neurons. Each datapoint represents technical replicate data from one 384-well plate well. Larger markers represent the medians each independent experiments. The boxplots show the median and the 2nd and 3rd quartiles of the data. Whiskers indicate 1.5X of the interquartile range. Welch’s unequal variances t-test was performed using the medians of independent differentiation experiments. ns = not significant, * = p < 0.05, ** = p < 0.01, *** = p < 0.001. (D) Scaled well and replicate median feature values aggregated into morphological profiles. Cosine similarity was used for clustering. (E) Top ten absolute mean differences between features of SNCA-4x and Control or SNCA-4x neurons treated with Prostratin. Means with standard deviations are shown. (F) Two first principal components (PCs) explaining most variation in the dataset after Principal Component Analysis (PCA). Each data point represents mean data from one well. (G) Absolute Pearson correlation of all features with the first ten PCs after PCA. Representative images illustrate cellular αSyn staining patterns. All data in this figure was generated in four independent experiments each containing at least 12 technical replicates per condition.

Next, we performed Principal Component Analysis (PCA) using the morphological profile data to visualize overall differences between compound-treated and untreated cells. The first two principal components (PCs) accounted for 57% of the variance and separated the tested cell lines and Prostratin-treated neurons, indicating significant differences between them (**Figure 1F**). The remaining PCs each accounted for 11% or less of the variance, suggesting lesser contribution to the overall variance (**Figure S1F**). To understand which features were mainly responsible for distinguishing the experimental categories, we calculated the absolute Pearson’s correlation between the first ten PCs and each morphological feature. PC1 correlated mostly with αSyn signal texture-related features, indicating a different αSyn subcellular distribution pattern in the neurons. PC2 and PC3 correlated with signal radial distribution and nucleus-related features, similar to the results we obtained when comparing experimental conditions pairwise using the mean absolute difference (**Figure 1G**, **Figure S1G**). Manual inspection of images confirmed these findings, showing grainy αSyn localization outside the nucleus in SNCA-4x cells (**Figure 1G**). This indicated that radial distribution and texture features indeed capture αSyn localization patterns. As a result, high-dimensional morphological profiling, by capturing a wide array of cellular characteristics, allowed us to identify subtle differences between experimental conditions that may not be apparent through visual inspection alone.

### Compound modulation shifts a SNCA-4x towards a control morphological profile

Based on our observation that small molecular compounds, such as Prostratin, can partially rescue SNCA-4x mDA neuron morphological profiles, we wondered whether more potent small molecules could be identified using a morphological profile screening approach. We assembled a 1,020 compound library which included 734 FDA-approved compounds, 435 of which can pass the blood-brain-barrier, 120 molecules targeting PD-relevant processes, and 166 diverse bioactive molecules (**Figure 2A**, **Table S3**). Cryopreserved D30 mDA neurons were cultured in 384-well plates until D33, treated with 5 μM of each compound for 72 hours and stained (**Figure 2B**). Each compound was tested in 4 replicates across 20 384-well plates, with 36 control wells per plate. Control conditions established baseline and maximum effect profiles (SNCA-4x + DMSO and SNCA-corr + DMSO, respectively). Morphological profiles were generated from 307,200 images and all data was aggregated per well (**Figure 2B**).

**Figure 2:**
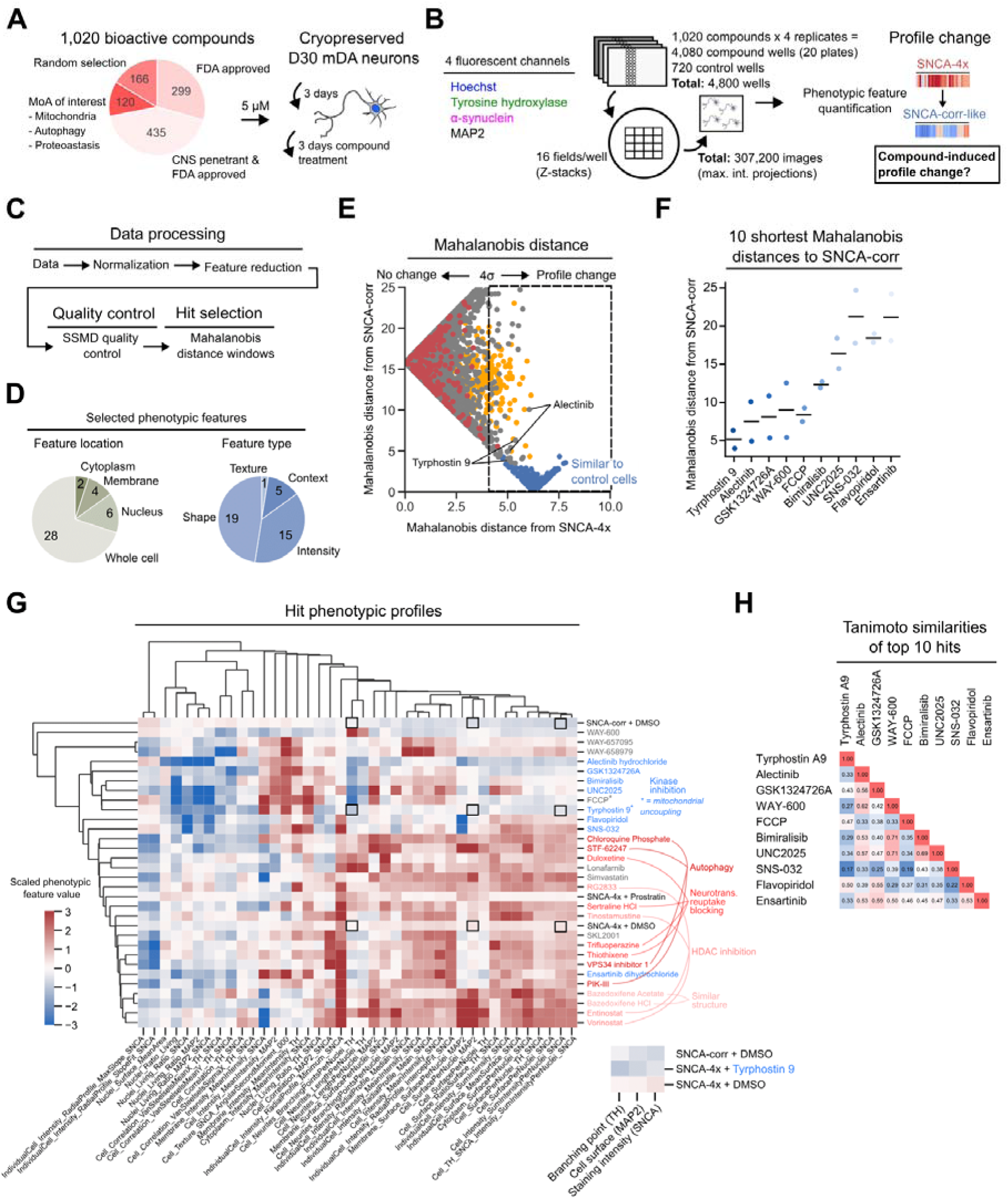
Compound modulation shifts a SNCA-4x towards a control morphological profile. **(A)** Overview about screening library composition and neuron treatment scheme. **(B)** Schematic overview of image data generation. Morphological features were calculated from imaging data to measure compound-induced morphological profile changes from a SNCA-4x towards a SNCA-corr-like profile. **(C)** Schematic of morphological profile data processing workflow. **(D)** Overview of morphological feature types used for hit selection after removal of redundant features using a Pearson correlation threshold of 0.95. **(E)** Morphological profile Mahalanobis distances of compound treated mDA neurons relative to SNCA-4x and SNCA-corr morphological profiles. The black dashed line indicates the selection window based on 4σ from the SNCA-4x median. Each datapoint represents mean data from one well and for compounds the mean value of all four replicates is shown. **(F)** Compounds inducing morphological profiles with the 10 shortest mean Mahalanobis distances to the median SNCA-corr morphological profile. **(G)** Selected hits clustered by the Cosine similarity of their respective morphological profiles. Compounds are color coded by attributed biological activity. (H) Tanimoto similarities of top 10 selected compounds.

We aimed to identify compounds shifting SNCA-4x profiles towards SNCA-corr-like phenotypes. Morphological profiles were Z-score normalized for comparability (**Figure 2C**, **Figure S2A**). Pearson’s correlation reduced the initial 127 features to 40 non-redundant ones, mostly describing cellular area, shape, and TH and αSyn staining intensities (**Figure 2D**, **Figure S2B**). Plate quality was assessed using the Strictly Standardized Mean Difference (SSMD) of the Mahalanobis distances between treatment groups. All plates showed significant differences (**Figure S2C**). Linear Discriminant Analysis (LDA) was used to compare each compound profile with the reference profiles. LDA achieved a classification accuracy F1 score of 0.96 in a test data set and 0.93 ± 0.04 in a 10-fold cross-validation, indicating a high ability to correctly predict whether compound-induced morphological profiles resemble one of the reference classes (**Figure S2D**). Most compounds had no effect on the SNCA-4x profile, while some induced SNCA-corr-like profiles (**Figure S2E**).

We aimed to identify compounds that could shift SNCA-4x profiles towards SNCA-corr-like phenotypes. Morphological profiles were compared using the Mahalanobis distance, a metric that measures how different two profile vectors are. We identified 28 compounds with a mean distance of at least 4 to the SNCA-4x median profile (**Figure 2E**, **Table S1**). Of these, the 10 compounds with the shortest distance to the SNCA-corr profile were considered to partially rescue the phenotype. Tyrphostin A9 and Alectinib were the most similar to the SNCA-corr profile (**Figure 2F**). To validate our findings, we used Cosine similarity, a metric that measures whether two vectors (or morphological profiles) are pointing in the same direction. Nine out of the 10 compounds with the shortest Mahalanobis distance to SNCA-corr grouped in the same cluster as SNCA-corr, supporting a distance and directional similarity between their morphological profiles (**Figure 2F**, **Figure 2G**). Although the top hits’ morphological profiles were highly similar among each other and compared to the SNCA-corr profile, their chemical structure-based Tanimoto similarities differed considerably (**Figure 2H**). We validated the effect of the identified compounds on cellular phenotypes using data from the Joint Undertaking in Morphological Profiling (JUMP) consortium, an effort to create compound-induced morphological profiles in U2OS cells by CellPainting ^30,31^. Also here, in a different cell type and using a different staining assay, we observed that compounds with similar mode of action induced similar morphological profiles (**Figure S2F**). This supports the notion that the identified compounds, while acting on different targets, might possibly acting on related pathways leading to a similar phenotypic outcome. Tyrphostin A9 was originally developed as an epidermal growth factor receptor (EGFR) inhibitor, but similar to other tyrphostins has been linked to mitochondrial membrane fragmentation, reduced OXPHOS, and potentially neuroprotection via reduction of ROS ^32–35^. Since mitochondrial defects play an important role in PD pathology, we decided to further characterize Tyrphostin A9’s molecular mechanism in mDA neurons ^3^.

### Tyrphostin A9 rescues the overall mDA neuron morphological profile in a dose dependent manner

To establish hit compound dose-response effects we focused on cell viability and αSyn level assessment as well as on morphological profiling using six different compound concentrations ranging from 0.25-20 μM and a treatment duration of 72 hours (**Figure S3**, **Figure S4)**. For Tyrphostin A9, we observed that higher compound concentrations in SNCA-4x neurons resulted in a higher resemblance to SNCA-corr neuron morphological features (**Figure 3A**). To quantify this observation, we calculated the Cosine distance of the compound treated SNCA-4x morphological profile to the SNCA-4x + DMSO and SNCA-corr morphological profiles as a function of compound concentration. Cosine distance can be defined as 1 – Cosine similarity. The intuition behind this is that if two vectors are perfectly the same then the similarity is 1 (angle=0) and thus, the distance is 0 (1-1=0). We found that increasing compound concentrations decreased the distance between SNCA-4x and SNCA-corr morphological profiles, therefore increasing their similarity (**Figure 3B**). The opposite is true when SNCA-4x compound treated neurons were compared to SNCA-4x DMSO-treated morphological profiles. Here, the Cosine distance increased with increasing compound concentration, reflecting the change of phenotype towards the SNCA-corr profile. Next, we investigated the effect of increasing compound concentrations on three single morphological features: αSyn intensity, TH intensity and MAP2 neurite length (**Figure 3C**). Tyrphostin A9 concentrations >1 μM decreased the nuclear and cytoplasmic αSyn staining intensity. Tyrphostin A9 also positively impacted TH staining intensity. At higher concentrations we observed little effects on MAP2 neurite length, indicating no acute toxicity.

**Figure 3:**
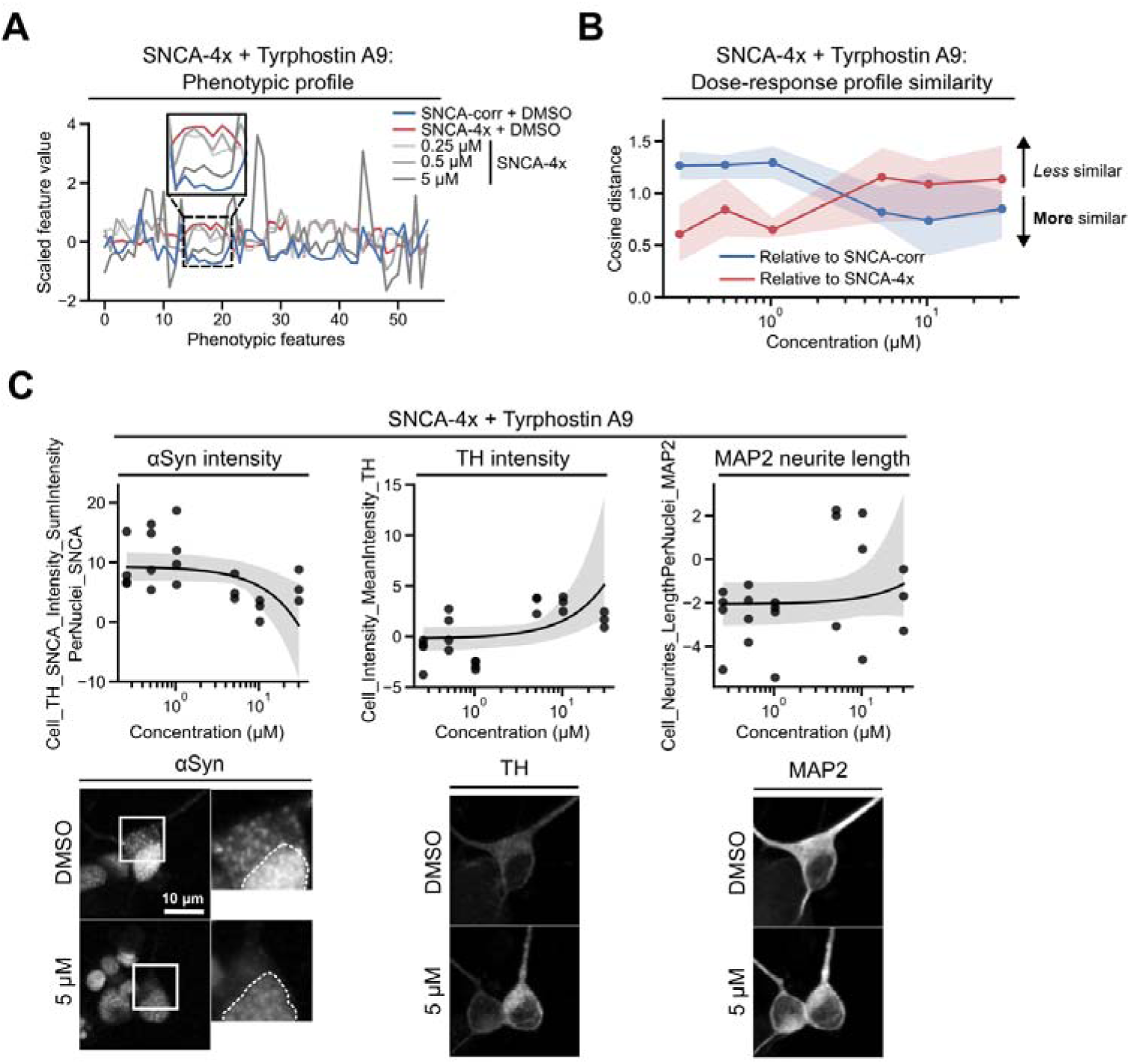
Tyrphostin A9 rescues the overall morphological profile in a dose dependent manner. **(A)** Line plot representation of morphological profiles of Tyrphostin A9 treated and untreated mDA neurons. D33 neurons were treated with 0.25-20 μM compound for 72 hours before fixation. **(B)** Cosine distances of SNCA-4x neurons as a function of compound concentration relative to DMSO-treated SNCA-4x and SNCA-corr neurons. **(C)** Dose-response curves for the three selected morphological features αSyn intensity, TH intensity, and MAP neurite length. Representative images of the same cell in each fluorescent channel are shown for DMSO and 5 μM compound treatment. Each compound concentration was tested at least in triplicate. Mean and 95% confidence interval are shown.

### Multiple identified hits induce mitochondrial uncoupling and ROS reduction

Tyrphostin A9 has been explored *in vitro* for its roles as neuroprotective agent, possibly functioning as an oxidative phosphorylation uncoupler reducing ROS ^32,33,36^. Mitochondrial uncoupling involves the decrease of the mitochondrial proton gradient disrupting the coupling between the electron transport chain and ATP synthesis. Whether Tyrphostin A9 can exert these effects in mDA neurons is currently unclear. We first assessed Tyrphostin A9’s ability to modulate the mitochondrial membrane potential using the JC-1 dye. We treated D33 mDA neurons for 72 hours with increasing concentrations of Tyrphostin A9 (0.01 µM to 10 µM) and observed a decrease in red fluorescence for concentrations >0.5 µM, indicating a concentration dependent decrease in mitochondrial membrane potential (**Figure 4A**). Valinomycin was used as positive control inducing a very low membrane potential. While FCCP and Tyrphostin A9 lowered the mitochondrial membrane potential at the lowest doses, also other molecules with morphological profiles similar to Tyrphostin A9, such as Tinostamustine or VPS34 inhibitor 1, led to a decreased mitochondrial membrane potential (**Figure S5**). Since a reduction in mitochondrial membrane potential is often associated with decreased ROS levels, we used a dye-based ROS assay to measure ROS levels in mDA neurons upon compound exposure. Tyrphostin A9, at concentrations >0.5 µM reduced ROS down to the level of the positive control antioxidant N-acetyl-cysteine (NAC) (**Figure 4B**). Interestingly, also multiple other molecules with morphological profiles similar to Tyrphostin A9, such as Simvastatin, Trifluoperazine, or WAY-600 lowered neuronal ROS levels (**Figure 2G**, **Figure S6**). We hypothesized that Tyrphostin A9 might lower mitochondrial membrane potential and ROS due to its function as protonophore facilitating proton transfer across membranes. We used a Rhod-FF assay which measures intracellular Ca^2+^ levels and found only small baseline differences between SNCA-corr and SNCA-4x mDA neurons, while the addition of Tyrphostin A9 to SNCA-4x mDA neurons increased intracellular Ca^2+^ levels by more than 70% (**Figure 4C**). Addition of the known protonophore Ionomycin further increased intracellular Ca^2+^ levels demonstrating that Tyrphostin A9 likely uncouples the mitochondrial membrane potential via facilitating the entry of protons, thereby decreasing oxidative phosphorylation and ROS levels. In neurons protonophores are also expected to disrupt the neuronal membrane potential by decreasing the proton gradient, leading to a decrease in the driving force for the generation of action potentials, which might negatively impact neuronal firing. Using a microelectrode array (MEA) we measured the neuronal firing rate before and after Tyrphostin A9 treatment (**Figure 4D**). Before treatment ca. 60-day old SNCA-corr mDA neurons showed approximately 2.5-fold higher firing rates than SNCA-4x neurons. When treating the mDA neurons for up to 72 hours with 2.5 μM Tyrphostin A9 we observed a sharp drop in firing already 30 minutes after treatment. After >24 hours of treatment a slight recovery effect was observed and firing rates began to increase mildly. Taken together, our findings suggest that the identified compounds, although affecting different targets, might be influencing related pathways, resulting in a similar phenotypic outcome such as mitochondrial membrane potential and ROS lowering.

**Figure 4:**
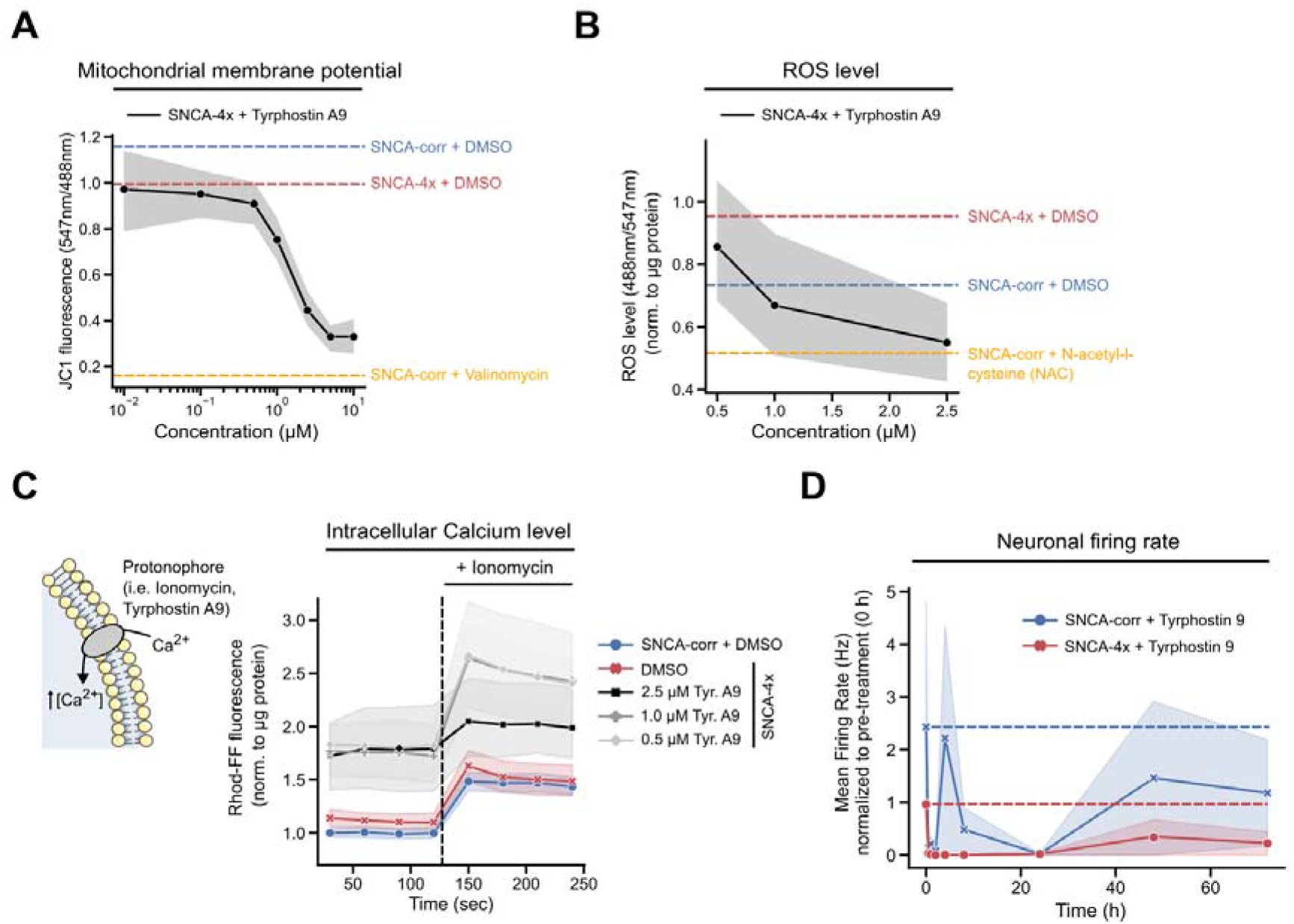
Tyrphostin A9 acts as ROS-reducing mitochondrial uncoupler in mDA neurons. (A) mDA neurons were treated with 0.1-10 μM Tyrphostin A9 for 72 hours and the mitochondrial membrane potential was measured using the JC1 dye. JC1 fluoresces red at high membrane potential and green at lower membrane potentials. Three independent experiments with at least three technical replicates each were performed. **(B)** mDA neurons were treated with 0.5, 1, and 2.5 μM Tyrphostin A9 for 72 hours and the ROS level was determined using the CellROX dye. The obtained signal was normalized to total protein content. Three independent experiments with at least three technical replicates each were performed. **(C)** *Left panel:* Scheme illustrating the protonophore mode of action. *Right panel:* mDA neurons were treated with 0.5, 1, and 2.5 μM Tyrphostin A9 for 72 hours and the intracellular Calcium level was measured using the Rhod-FF dye for 250 seconds before and after the addition of Ionomycin. The obtained signal was normalized to total protein content. Two independent experiments with at least three technical replicates each were performed. **(D)** Neuronal activity was measured in ca. 60-day old mDA neurons using a microelectrode array (MEA). The neuronal firing rate was assessed before and up to 72 hours after treatment with 2.5 μM Tyrphostin A9. Neuronal activity was measured in two independent experiments with at least three technical replicates each. For all data, the mean and 95% confidence interval is shown.

### The Tyrphostin A9 structural analogue AG879 induces mitochondrial uncoupling

Tyrphostins represent a large family of small molecules with various attributed neuroprotective modes of action. Next to their role as mitochondrial uncouplers, tyrphostins have been described as antioxidants or as having a positive effect on cellular glutathione levels ^33^. We therefore compared a range of Tyrphostin A9 structural analogues using various biochemical assays to establish a Tyrphostin A9 structure-function relationship in mDA neurons (**Figure 5A**). FCCP is a potent mitochondrial uncoupler used during oxygen consumption rate (OCR) measurements and facilitates proton transport across mitochondrial membranes. This action collapses the proton gradient and disrupts the mitochondrial membrane potential. As a result, the electron flow through the Electron Transport Chain (ETC) is uninhibited, allowing oxygen consumption by complex IV to reach its maximum. The resulting OCR increase is used as an estimation of the maximal respiratory capacity of the cell (**Figure 5B**, left panel). Next to its use as a control compound, we also detected FCCP as one of the top hits during morphological profiling (**Figure 2F**). We therefore compared Tyrphostin A9 and its structural analogues to FCCP using an OCR assay. We found that, similar to FCCP, Tyrphostin A9 and its closest structural analogues AG1024 and AG879 led to a strong increase in maximal respiratory capacity, highlighting their mitochondrial uncoupling activity (**Figure 5B**, right panel). Next, we determined compound cytotoxicity by measuring Lactate dehydrogenase (LDH) in the culture medium. LDH is a stable enzyme, present in all cell types and rapidly released into the culture medium upon damage of the plasma membrane. None of the tested compounds induced significantly more cell death than the DMSO control when used at 0.5-2.5 μM for 72 hours. Cell lysate was used as control corresponding to the maximum possible amount of released LDH (**Figure 5C**). JC1 dye fluorescence was measured to determine the mitochondrial membrane potential. Tyrphostin A9 and its most closely related structural analogue Tyrphostin AG879 decreased the membrane potential as measured by the 547nm/488nm fluorescence ratio from 0.95 to approximately 0.2. The second most similar analogue Tyrphostin AG1024 also decreased the membrane potential, although to a lesser extent (**Figure 5D**). In line with the decreased membrane potential Tyrphostins A9 and AG879 also reduced ROS levels to ca. 80% of the DMSO baseline level (**Figure 5E**). Next, we assessed glutathione levels. In healthy cells, both reduced glutathione (GSH) and its oxidized form (GSSG) are present, with the reduced form (GSH) making up the majority, while the oxidized form (GSSG) increases during oxidative stress. Although we expected an GSH increase due to ROS reduction, we observed that Tyrphostins A9, AG879 and AG1024 decreased GSH ratio, possibly due to lower overall glutathione synthesis after ROS decrease. Tyrphostin A9 lead to the strongest decrease of 48% at 2.5 μM (**Figure 5F**). Mitochondrial uncoupling also strongly reduces cellular ATP production. We therefore tested the effects of all compounds on mDA neuron ATP level. We found that the compounds with the strongest uncoupling effects (Tyrphostins A9, AG1024, and AG879) reduced ATP levels the most (**Figure S7A**). mDA neuron treatment with the mitochondrial complex I inhibitor Rotenone led to similarly low ATP levels as Tyrphostin A9 treatment alone, indicating a near maximal Tyrphostin A9 uncoupling effect (**Figure S7B**). Since close Tyrphostin A9 structural analogues such as AG879 and AG1024 reproduce its cellular effects, the shared structural features may be critical for the molecule’s function in mDA neurons.

**Figure 5:**
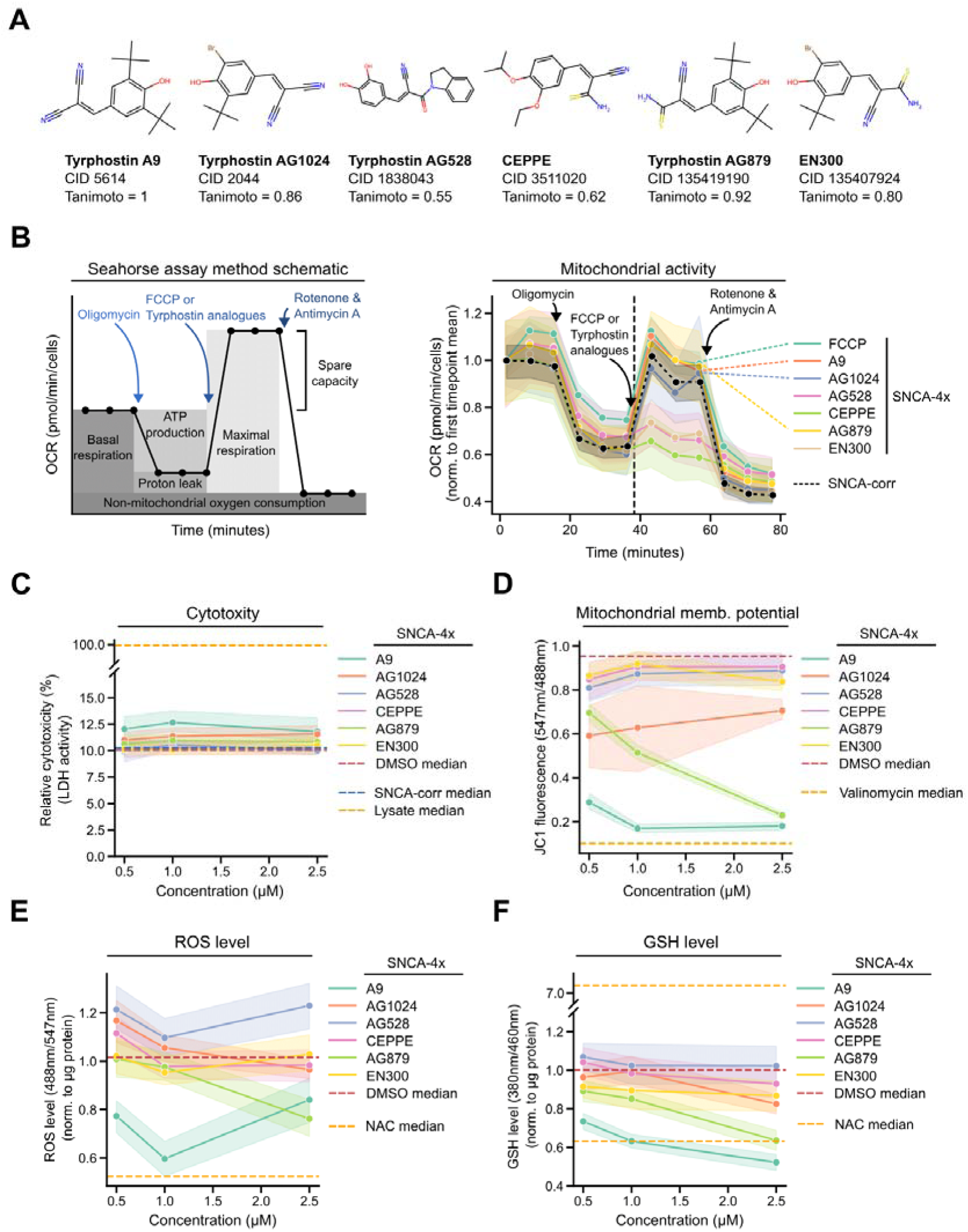
Tyrphostin A9 structural analogues reduce mitochondrial membrane potential and ROS. **(A)** Molecular structures of Tyrphostin A9 and five selected structural analogues including their names, Compound ID (CID), and Tanimoto similarity coefficient relative to Tyrphostin A9. **(B)** Oxygen consumption rate (OCR), a proxy readout for mitochondrial respiration, was measured in D30 mDA neurons treated with 2.5 μM compound. (C-F) D33 mDA neurons were treated with 0.5-2.5 μM of each compound for 72 hours. **(C)** Lactate dehydrogenase (LDH) activity was measured in the culture medium to determine relative cytotoxicity compared to cellular lysate (=100%). **(D)** Mitochondrial membrane potential was measured using the JC-1 dye. **(E)** ROS level was determined using the ROS-ID detection kit and the obtained signal was normalized to total protein content. **(F)** Reduced glutathione (GSH) was measured using monochlorobimane (MCB) and the obtained signal was normalized to total protein content. Three independent experiments with at least three technical replicates were performed. The mean and 95% confidence interval are shown. Dashed lines indicate the median values of the respective controls.

### Tyrphostin A9 induces αSyn lowering

Our phenotypic compound screen revealed that mDA neuron αSyn levels were lowered by Tyrphostin A9 treatment (**Figure 3C**). Since immunofluorescent staining is not sensitive enough to differentiate between αSyn monomers, oligomers, and larger structures such as fibrils, we performed Western blotting of Tyrphostin A9 treated and untreated >D50 mDA neuron lysates (**Figure 6A**). We observed a 2-fold reduction of both low molecular weight αSyn (≈15 kDa) as well as higher molecular weight αSyn species. Most cellular proteins are selectively degraded by the proteasome via ubiquitination. Caspase and chymotrypsin are part of the proteasomal protein degradation machinery, contributing to the proteolytic activities of the proteasome. Using plate reader assays we therefore checked how caspase and chymotrypsin activities are impacted by Tyrphostin A9 in mDA neurons. Both enzymes’ activities were not significantly different between SNCA-corr and SNCA-4x mDA neurons, while addition of Tyrphostin A9 rather decreased (≈ 50%) or did not impact caspase- or chymotrypsin-related proteasomal degradation, respectively (**Figure 6B**). As expected, the known proteasome inhibitor MG132 strongly decreased both caspase and chymotrypsin activities in mDA neurons by >70%. Since Tyrphostin A9 does not positively impact proteasomal protein degradation in mDA neurons, the observed lower αSyn levels after compound treatment might result from an increased alternative protein degradation pathway.

**Figure 6:**
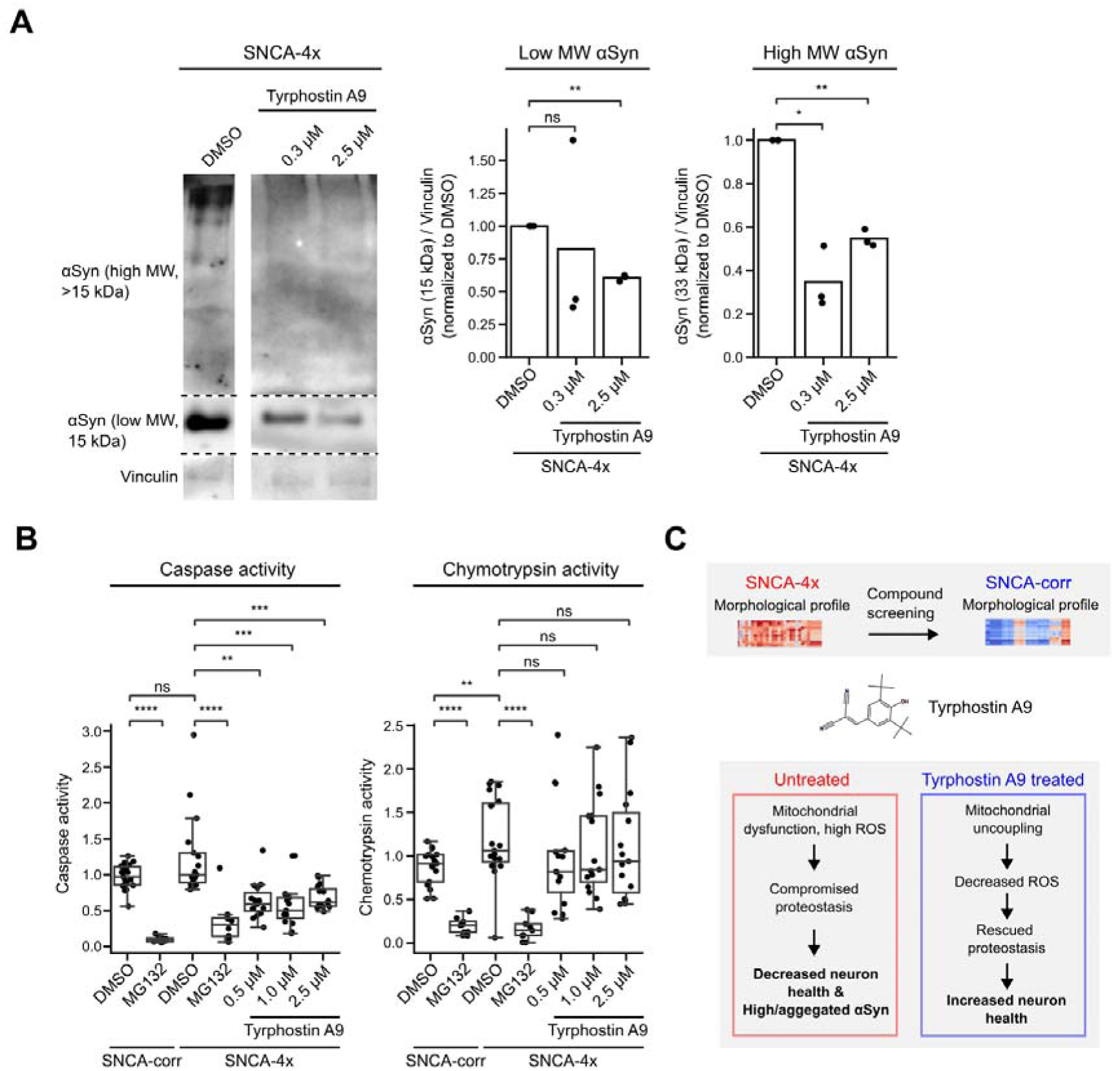
Tyrphostin A9 decreases αSyn protein expression. **(A)** Anti-αSyn Western blotting of mDA neuron lysates. Before lysis >D50 mDA neurons were treated with 0.3 or 2.5 μM Tyrphostin A9 for 72 hours. Three independent neuronal differentiations were performed. Gel images are cropped. Full-length blots/gels are presented in Figure S8. **(B)** Caspase and Chemtrypsin activities were determined in approximately D30 mDA neurons treated with 1 or 2.5 μM Tyrphostin A9 or the proteasome inhibitor MG132 for 72 hours. Three independent experiments with at least three technical replicates each were performed. The boxplots show the median and the 2nd and 3rd quartiles of the data. Whiskers indicate 1.5X of the interquartile range. Welch’s unequal variances t-test was performed using the technical replicates of all independent experiments. ns = not significant, * = p < 0.05, ** = p < 0.01, *** = p < 0.001. **(C)** Scheme depicting the use of morphological profiling to identify compounds that produce a healthy control-like (SNCA-corr) profile in mutant (SNCA-4x) mDA neurons. The identified compound Tyrphostin A9 increases mitochondrial uncoupling and decreases ROS and αSyn levels which can explain the rescued morphological profile.

## Discussion

We used microscopy-based morphological profiling to characterize the effects of 1,020 FDA-approved or PD pathobiology-related small molecules on the overall morphological appearance of human iPSC-derived SNCA-4x mDA neurons. We scored compounds based on their potential to induce a morphological profile resembling SNCA-corr mDA neurons (**Figure 2**). Phenotypic or morphological profiling is increasingly used in iPSC-derived cells, primary cells or brain slice systems to study neuronal diversity and to complement single cell RNA sequencing data ^37–40^. Previously, we established morphological profiling in LRRK2 G2019S mutated iPSC-derived mDA neurons and showed that multiple LRRK2 inhibitors can partially rescue the neuronal morphological profile ^8^. Drug screens in mDA neuron model systems have been performed before, however either using only small screening libraries, non-human cell systems such as zebrafish, or very young mDA neuron progenitors (<20 days of differentiation) ^41–44^. This study combines detailed morphological profiling in human mDA neurons with a compound screen using a larger library (>1,000 compounds). Although morphological profiling has the potential to identify completely novel targets, a limitation is that it typically provides little information about the actual potential compound targets, requiring detailed target and mode of action deconvolution studies.

In our study, we focused on mitochondrial uncoupling since two of our top scoring compounds (FCCP, Tyrphostin A9) are annotated as protonophores and mitochondrial uncouplers. Originally developed as an EGFR inhibitor, Tyrphostin A9 stood out due to its αSyn reducing and TH increasing effects in our morphological profiling assay as well as the molecule’s previously described role in neuroprotection (**Figure 3**) ^33–35^. We found that the molecule promotes mitochondrial uncoupling and reduces ROS in mDA neurons in line with its previously described protonophore properties in other cell types (**Figure 4**) ^33,36,45^. Tyrphostins have been described as mitochondrial uncouplers for almost half a century, while mitochondrial uncoupling as a neuroprotective strategy is more recent, but has faced several safety-related hurdles during pre-clinical drug development ^46–49^. In mDA neurons we made the observation that next to its mitochondrial uncoupling effect, Tyrphostin A9 also decreased neuronal activity which relies on charge-dependent action potentials (**Figure 4D**). Tyrphostin A9 did not permanently damage the neurons, indicated by the gradual partial activity recovery over time, intact morphology, and low extracellular LDH levels (**Figure 3**, **Figure 4D**, **Figure 5C**). Given that pace making is a critical physiological function of mDA neurons in the *substantia nigra*, this observation raises concerns about potential detrimental effects of Tyrphostin A9. Importantly, we do not claim that Tyrphostin A9 itself holds therapeutic potential. However, here we show that morphological profiling in a PD-relevant human model system is able to detect compound classes with potentially undervalued neuroprotective modes of action. Future research should address the safety of mitochondrial uncoupling as a neuroprotective strategy and explore the link between Tyrphostin A9-induced EGFR inhibition and neuroprotection. Recently the blocking of mitochondrial complex I in pro-inflammatory microglia has been shown to protect the central nervous system against neurotoxic damage in mice ^50^. An attractive future neuroprotective therapeutic strategy involving mitochondrial uncouplers might therefore involve non-cell autonomous mechanisms. The molecular characterization of non-cell autonomous mechanisms in neurodegeneration is becoming increasingly feasible by using more complex iPSC-derived model system involving multiple central nervous system cell types in the form of co-culture experiments or organoids ^51–53^.

A surprising observation we made is Tyrphostin A9’s ability to reduce the low and high molecular weight αSyn levels in mDA neurons by about 50% (**Figure 3C**, **Figure 6**). We excluded a direct effect on caspase or chymotrypsin related proteasomal degradation mechanisms since Tyrphostin A9 either did not impact or even decreased these modes of protein degradation. Except for one study describing a role of Tyrphostin AG1024 in accelerating the degradation of Retinoblastoma Protein (pRb) likely involving the proteasome, tyrphostins have not been directly linked to enhanced protein degradation ^54^. Next to proteasomal protein degradation, the involvement of lysosomes in Parkinson’s disease progression through two major autophagic pathways, chaperone-mediated autophagy and macroautophagy, and their central role in degrading αSyn is well documented ^2^. A single study demonstrated that Tyrphostin AG490, via JAK2 inhibition, prevented the hormone-receptor complex Epo-EpoR from targeting lysosomes in the UT-7 human cell line, highlighting a possible link between tyrphostin action and lysosomal biology ^55^. Although we did not elucidate the exact sequence of events in our mDA neuron model, oxidative stress can cause proteins to misfold, leading to proteostatic stress. Conversely, misfolded proteins can disrupt mitochondrial function, increasing ROS production and oxidative stress, leading to dopamine oxidation ^3^. αSyn could play a key role in the harmful effects of oxidized dopamine. It has been shown that oxidized dopamine can alter and stabilize αSyn protofibrils *in vitro*, and an increased expression of αSyn in SH-SY5Y cells heightened their susceptibility to an excess of dopamine ^56,57^. In PD patient-derived iPSC neurons, a rise in oxidized dopamine and a reduction in αSyn solubility has been observed ^26^. Moreover, mitochondrial uncouplers can induce mitophagy which in turn could decrease the α-synuclein that is accumulated and aggregates in mitochondria ^58–60^. Future work will need to establish the molecular connection between mitochondrial uncoupling and the increased αSyn degradation which we observed.

## Material and Methods

### Origin of iPSC lines and differentiation into mDA neurons

*SNCA* iPSC lines (originally named AST23 and AST23-2KO) were provided and generated by Dr. Tilo Kunath from The University of Edinburgh. The parental cell line AST23 was first described in Devine *et al.* ^19^. The generation of the SNCA-corr cell line AST23-2KO is described in Chen *et al.* ^61^. Both cell lines are deposited as EDi001-A and EDi001-A-5 in the hPSCreg repository, respectively (https://hpscreg.eu). iPSCs were cultured on Geltrex-coated (Thermo Fisher Scientific) dishes in StemMACS iPS-Brew XF (Miltenyi Biotech). The medium was changed daily, and cells were passaged twice a week using 0.5 mM EDTA in PBS (Thermo Fisher Scientific). Mycoplasma testing was performed twice per month. mDA neurons were differentiated using a modified protocol based on Kriks *et al.* ^62–64^. In summary, iPSCs were initially placed on Geltrex-coated six-well plates or T75 flasks at a density of 2×10^5^ cells/cm2 in StemMACS iPS-Brew XF with 10 μM Y-27632 (Hiss). The following day, the medium was changed to KnockOut DMEM medium with KnockOut serum replacement (both from Thermo Fisher Scientific), supplemented with 200 nM LDN19318 (Axon Medchem) and 10 μM SB431542 (Biozol) for dual SMAD-inhibition. On the second day, 100 ng/mL Shh C24II (Miltenyi Biotech), 2 μM Purmorphamine (Miltenyi Biotec), 100 ng/mL FGF8 (Peprotech), and 3 μM CHIR99021 (Miltenyi Biotec) were also added to the medium. After a period of 5 days, the medium was gradually transitioned to Neurobasal medium (Thermo Fisher Scientific), and SB431542 was removed from the medium. Starting from day 7, cells were cultivated only in the presence of LDN19318 and CHIR99021. On day 11, cells were transferred to Neurobasal/B27/L-glutamine medium with only CHIR99021. On day 13, cells were moved onto Geltrex-coated dishes in Neurobasal/B27/L-glutamine medium with supplements of 20 ng/mL BDNF, 20 ng/mL GDNF (both from Cell Guidance Sys.), 220 μM L-ascorbic-acid (Sigma-Aldrich), 10 μM DAPT (Axon Medchem), 1 ng/mL TGF-βIII (Peprotech), 0.5 mM dibutyryl-cAMP (Enzo Life Sciences), and 10 μM Y-27632 (Hiss). Cells were kept in the same medium but without Y-27632. Around days 23–25, cells were separated using StemPro Accutase (Thermo Fisher Scientific) and placed at a density of 1.4×10^5^ cells/cm2 onto Geltrex-coated dishes. To remove non-neuronal cells, cultures were exposed to 1 μg/mL Mitomycin C for 2 hours on day 26. On day 30, neuronal cultures were separated using StemPro Accutase with 10 μM Y-27632 and singularized. Cells were counted and preserved at 2.5×10^6^ cells/vial in CryoStor CS 10 tubes (Sigma-Aldrich).

### Neuronal culture

30 days old mDA neurons were thawed and spun (400 g, 5 min, RT) in a basal medium (**Table S2**) supplemented with a ROCK inhibitor (Tocris #1254). The cell pellets were then reconstituted in a differentiation medium (**Table S2**) also supplemented with a ROCK inhibitor. 384-well plates (Perkin Elmer, Phenoplate poly-D-lysine-coated #6057500) were prepared with a coating of 10 μg/ml laminin at 4°C overnight. Using trypan blue (Invitrogen, #T10282) and a Countess automated cell counter (Invitrogen), cells were seeded in the 384-well plates at a density of 10×10^3^ cells/well. Edge wells were not used for seeding and were filled with PBS instead. Typically, the thawed cells were incubated at 37°C and 5% CO2 for 6 days until they reached day 36, with changes in the differentiation medium occurring at D33. The processes of plate coating, cell seeding, and medium changes were automated using an Bravo Automated Liquid Handling Platform with 384ST head (Agilent) and a Multidrop Combi Dispenser (ThermoFischer Scientific).

### Compound library and compound treatment

1,020 small molecules were selected from a bioactive compound library (Selleckchem), based on PD-relevant pathobiology, FDA approval status, and blood-brain-barrier penetrance (**Table S3**). *Prostratin treatment:* Neurons were cultured until D33, treated with 5 μM Prostratin (Biotechne) or DMSO for 72 hours and immunofluorescently stained. *Primary screening:* Cryopreserved mDA D30 neurons were cultured in 384-well plates until D33, treated with each compound at a concentration of 5 μM for a duration of 72 hours, and subsequently stained. Each compound underwent testing in four technical replicates. Compound wells were randomly assigned between technical replicates. 36 wells per plate were designated as control wells. The control conditions were used to establish the baseline and maximum effect profiles, specifically with SNCA-4x + DMSO serving as the baseline and SNCA-corr + DMSO as the maximum effect profile. Compounds and DMSO were dispensed using a Bravo Automated Liquid Handling Platform with 384ST head (Agilent). *Dose-response testing:* Identical to primary screening but with six different compound concentrations ranging from 0.25-20 μM. *Biochemical assays:* >D30 neurons were treated with 0.5-2.5 μM compound for 72 hours. *Microelectrode array (MEA):* Ca. D60-old mDA neurons were treated up to 72 hours with 2.5 μM Tyrphostin A9. *Western blotting:* >D50 mDA neurons were treated with 0.3 and 2.5 μM Tyrphostin A9 for 5 days.

### Immunofluorescent staining

Neurons were fixed using 4% PFA (EMS Euromedex, #15710) for a duration of 20 minutes, followed by three PBS washes (Gibco, #14190136). The cells were then permeabilized and blocked using a solution of 10% FBS (Sigma, #F7524) and 0.1% Triton X-100 (Sigma, #T9284) in PBS for an hour. Primary antibodies against TH, αSyn, and MAP2 (**Table S2**) were prepared in an antibody dilution buffer (PBS with 5% FBS and 0.1% Triton X-100) and left to incubate with the cells at 4°C overnight. This was followed by three PBS washes. Secondary antibodies and Hoechst (**Table S2**) were added to the cells in the antibody dilution buffer and left for an hour at room temperature, followed by another three PBS washes. Immunofluorescent staining was automated using an EL406 plate washer and dispenser (Biotek).

### Image acquisition and feature extraction

Imaging was performed on a Yokogawa CV7000 microscope in scanning confocal mode using a dual Nipkow disk and a cooled sCMOS camera with 2,560×2,160 pixels and a pixel size of 6.5 μm without pixel binning. The system’s CellVoyager software, the 405/488/561/640-nm solid laser lines, and a dry 40X objective were used to acquire 16 images in a 4×4 orientation from three 1 µm-separated Z-planes from the center of each well. Wells were visited in a row-wise “zigzag” fashion. Maximum intensity projections were performed. Image segmentation and feature extraction was performed with our in-house software PhenoLink (https://github.com/Ksilink/PhenoLink). Image segmentation was performed on 4×4 stitched and illumination corrected images. Using intensity and size thresholds specific to each of the four imaging channels (Hoechst, TH, αSyn, and MAP2) we extracted a total of 124 morphological features, including count, texture, context, intensity and shape features (**Table S4**). We used location constraints to identify subcellular compartments. For example, by subtracting the segmented nucleus from the whole cell we obtained the cytoplasm or by calculating a 4 pixel-wide area around the outer edge of the cell we estimated the location of the cell membrane.

### Data processing & morphological profile generation

All data manipulation and transformation processes were carried out using Python’s Pandas package ^65^. All plots were generated using the Seaborn package ^66^. All data processing and plotting can be reproduced using the provided Python Jupyter Notebooks and accompanying raw data. Briefly, raw morphological profiles were created per well by using the information from all measured 127 morphological features (**Table S4**). First, per feature a mean feature value from all acquired 16 image fields per well was calculated. The feature values were subjected to a standard scaling process using the StandardScaler function from the sklearn.preprocessing module in Python, resulting in the creation of 127-feature raw morphological profiles per well ^67^. To determine the most defining morphological features per treatment condition, the morphological profiles were grouped based on their experimental replicates and treatments. The absolute differences in the mean values of the grouped data were calculated and followed by the computation of the mean and standard deviation of these absolute differences. We employed Principal Component Analysis (PCA) to compare morphological feature profiles across biological conditions and treatments. PCA was performed with 10 components and the explained variance ratio for each of the principal components was calculated. To measure how each original feature contributed to the principal components, the correlation between the original features and the first 10 principal components was calculated.

### Primary screening data processing and hit selection

The raw data was composed out of 307,200 images from 4,800 wells distributed over 20 384-well plates. For all fluorescent channel images (Hoechst, TH, αSyn, and MAP2) we performed image segmentation and feature extraction, giving rise to a raw 127-feature morphological profile. Per plate, extreme outlier features were eliminated using a robust standard deviation-based method. This involved calculating the absolute difference between each data point and the feature median, and retaining data that was within 4σ of the Median Absolute Deviation (MAD). Next, data was normalized to allow comparisons between plates. We applied a robust Z-score normalization, during which the original data was adjusted based on its deviation from the median of the DMSO control data and was scaled by the MAD of the DMSO control data. This method is robust to outliers as it uses median and MAD, which are less sensitive to extreme values. Feature reduction was performed in two separate steps. Since the morphological profile contains an overrepresentation of texture features, first the two most distinguishing texture features were selected by calculating the z-factors for each feature, sorting them in descending order, and picking the top two. In the second step, features with less than 95% pairwise Pearson correlation were retained. In total, 40 features remained (**Figure S2B**). Plate quality was assessed using the Strictly Standardized Mean Difference (SSMD) based on the 40-feature Mahalanobis distance of SNCA-4x + Prostratin-treated control wells to SNCA-4x + DMSO control wells. SSMD values for all 20 plates were <-1, indicating a good plate quality. We then calculated the 40-feature Mahalanobis distances of all SNCA-4x compound-treated wells to the SNCA-4x + DMSO and SNCA-corr + DMSO control wells. We applied a Mahalanobis distance threshold of >4 to identify compounds inducing morphological profiles different from SNCA-4x + DMSO control profiles. Additionally, we selected compounds based on close Mahalanobis distances to SNCA-corr + DMSO control profiles, thereby selecting compounds that induce morphological profiles similar to healthy control neurons. All data processing and hit selection steps can be reproduced by using the provided Python Jupyter Notebooks.

### Evaluation of mitochondrial membrane potential

Mitochondrial membrane potential (Δψm) was estimated using the fluorescent probe JC-1 (ThermoFisher, T3168). Briefly, 60×10^3^ mDA neurons were seeded per well in a Geltrex-coated black, clear-bottom 96-well cell culture plate and treated with selected compounds for three days. On neuronal maturation day 33, the neuronal medium was removed and cells were incubated with 100μL/well of fresh neuronal medium containing 10 μΜ of JC-1 dye for 30 minutes at 37°C. After incubation, cells were washed three times with pre-warmed DPBS (ThermoFisher, #14190094). Fluorescence was measured using a Cytation 5 (Bio-Tek) fluorescence plate reader, with excitation at 560 nm and emission at 610 nm for the aggregated JC-1 and excitation at 500 nm and emission at 550 nm for the monomeric JC-1. Mitochondrial membrane potential was estimated by calculating the ratio of aggregated to monomeric JC-1 fluorescent signal. Statistical analyses were conducted on data from, at least, three independent biological replicates.

### Measurement of intracellular ROS and glutathione levels

Intracellular ROS levels were measured using the ROS-ID Total ROS/Superoxide detection kit (Enzo, Enz-51010), following the manufacturer’s instructions. Briefly, 60×10^3^ mDA neurons were seeded per well in a Geltrex-coated black, clear bottom 96-well cell culture plate. Neurons were treated for three days with the selected compounds. On neuronal maturation day 33, the neuronal medium was removed and the cells were washed once with pre-warmed DPBS (ThermoFisher, #14190094). Cells were then incubated with 100 μL/well of ROS Detection Solution for 1 hour at 37°C. Finally, fluorescence is measured without removing the detection solution, using a Cytation 5 (Bio-Tek) fluorescence plate reader (bottom reading at Ex=490 nm, Em=520 nm). Fluorescence intensity was normalized to total cell protein levels, using the BCA Protein Detection Kit (ThermoFisher, #23227). Statistical analyses were conducted on data from three independent biological replicates. Simultaneously, intracellular GSH levels were measured using the monochlorobimane (MCB, AAT Bioquest, #76421-73-3) assay. In this procedure, 60×10^3^ mDA neurons were seeded per well in a Geltrex-coated black, clear-bottom 96-well cell culture plate and treated with selected compounds for three days. On neuronal maturation day 33, the neuronal medium was removed, and the cells were washed once with pre-warmed DPBS (ThermoFisher, #14190094). The cells were then incubated with 100 μL per well of a detection solution containing 50 μM MCB for 1 hour a, with excitation at 360 nm and emission at 480 nm t 37°C. After incubation, fluorescence was measured directly without removing the detection solution using a Cytation 5 (Bio-Tek) fluorescence plate reader, with excitation at 360 nm and emission at 480 nm. The fluorescence intensity readings were normalized to total cell protein levels, which were determined using the BCA Protein Detection Kit (ThermoFisher, #23227). Statistical analyses were conducted on data from three independent biological replicates.

### Evaluation of cytotoxicity

Cytotoxicity was estimated using the lactate dehydrogenase (LDH) assay (Canvax, CA0020), following the manufacturer’s instructions. 60×10^3^ mDA neurons were seeded per well in a Geltrex-coated black, clear bottom 96-well cell culture plate. Neurons were treated for three days with the selected compounds. On neuronal maturation day 33, 50 μL of media was obtained from each well and transferred to a 96-well cell culture plate, in order to evaluate the release of LDH into the cell culture media. It was added to an equal volume (50 μL) of mixed LSH assay buffer and dye solution and was incubated for 30 minutes, at room temperature, protected from light. Finally, 50 μL Stop Solution was added per well and the absorbance (OD) was measured using a Cytation 5 (Bio-Tek) plate reader (450 nm). LDH absorbance was normalized to total cell protein levels, using the BCA Protein Detection Kit (ThermoFisher, #23227). Statistical analyses were conducted on data from three independent biological replicates.

### Measurement of intracellular ATP levels

Intracellular ATP levels were evaluated using the CellTiter-Glo 2.0 (Promega, G9241), according to the product’s instructions. Briefly, 70×10^3^ mDA neurons were seeded per well in a Geltrex-coated black, clear-bottom 96-well cell culture plate and treated with selected drugs or compounds for three days. On neuronal maturation day 33, the neuronal medium was removed, and cells were washed once pre-warmed DPBS (ThermoFisher, #14190094). Cells were incubated with 75 μL per well of HBSS and an equal volume of the CellTiter-Glo 2.0 mix was added in each well. The plates were shaken for 2 minutes in RT, followed by 10 minutes of incubation in RT. Luminescence was measured using a Cytation 5 (Bio-Tek) plate reader. Luminescence intensity was normalized to total cell protein levels, using the BCA Protein Detection Kit (ThermoFisher, #23227). Statistical analyses were conducted on data from three independent biological replicates.

### Analysis of mitochondrial respiration

Oxygen consumption rates (OCRs) were measured using the Seahorse XFe96 FluxPak (Agilent, #102416-100) and the XF Cell Mito Stress Test Kit (Agilent, #103015-100), following the manufacturer’s instructions. For this experiment, 70×10^3^ mDA neurons were seeded per well in a Geltrex-coated 96-well Seahorse cell culture plate. On neuronal maturation day 33, the neuronal medium in each well was replaced with 175 µl of Seahorse XF DMEM Medium (pH 7.4, Agilent, #103575-100) containing 25 mM Glucose, 2 mM L-Glutamine, and 1 mM Sodium Pyruvate. The cells were incubated for 1 hour at 37°C without CO2. Meanwhile, the sensor cartridge, pre-equilibrated overnight in XF Calibrant solution at 37°C without CO2, was prepared with standard mitochondrial inhibitors: Oligomycin (2.5 µM), FCCP or tyrphostin analogs (2.5 µM), and a mixture of Rotenone and Antimycin A (1 µM each). After a further 30-minute incubation at 37°C without CO2, the sensor cartridge and the cell culture plate were loaded into the XFe96 Extracellular Flux Analyzer (Agilent). OCR was measured three times for basal respiration and after each injection of mitochondrial toxins. Post-assay, the cell culture medium was removed, and total protein content was determined using a BCA Protein Assay Kit (ThermoFisher, #23227). The OCR data were analyzed with XFe Wave Pro software (Agilent), normalized to total cell protein levels (based on the BCA assay), and exported using the Seahorse XF Cell Mito Stress Test Report Generator. Statistical analyses were conducted on data from three independent biological replicates. The study calculated various parameters related to oxidative phosphorylation, including basal respiration, proton leakage across the inner mitochondrial membrane, ATP-coupled respiration, maximal respiration, and spare respiratory capacity.

### Multi-electrode array (MEA)

Cryopreserved 30 days old neurons were thawed in a water bath and centrifuged (400g, 5 min, RT) in basal medium (Table S2) supplemented with ROCK inhibitor (Tocris, #1254). Cell pellets were resuspended in differentiation medium (Table S2) supplemented with ROCK inhibitor. 48-well CytoView MEA plates (Axion Biosystems, #M768-tMEA-48W) were coated with 15 μg/ml Polyethyleneimine solution (Sigma, #P3143) for 1 hour at 37°C followed by 10 μg/ml Laminin coating for 2 hours at 37°C. Using Tryphan Blue (Sigma, # T8154-20ML) and a Countess automated cell counter (Invitrogen), 80×10^3^ cells/well were seeded in the CytoView plate. Cells were incubated at 37°C and 5% CO_2_ for 26-33 days with differentiation medium changes twice a week. Once a week, 1μg/ml Laminin was added to the fresh media to maintain cell attachment. Neurons were then treated with compound for 72 hours. The electrical activity was recorded on the Maestro Pro multiwell MEA (Axion Biosystems) using the AxIS Navigator software (version 3.5.1, Axion Biosystems). Electrical activity was recorded for 10 minutes at a 12.5 kHz sampling rate and a 5.5 standard deviation threshold level for action potential detection. Before plate loading the device was allowed to equilibrate for approximately 30 minutes to 37°C and 5% CO2. Data analyses were performed using the Neural Metric Tool (version 3.1.7, Axion Biosystems). Activity is expressed as the mean firing rate across electrodes in a well.

### SDS gel electrophoresis and Western blotting

1.5×10^6^ mDA neurons were cultured for at least 50 days in Geltrex-coated 6-well plates with neuronal media and treated with compound for five days. At day 50, the neuronal media was removed and cells were washed with pre-warmed DPBS. DPBS was removed and cells were crosslinked by adding 1 mL pre-warmed PBSx1 supplemented with 2 mM disuccinimidyl glutarate (DSG, ThermoFisher, #20593) and protease inhibitors for 30 minutes at 37°C. Crosslinking was terminated by adding 50 mM Tris-HCl, pH=7.5 for 15 minutes at room temperature. Cells were collected and centrifuged for 10 minutes at 1000xg at room temperature. The supernatant was removed and the cell pellet was lysed using lysis buffer (100 mM NaCl, 5 mM EDTA, 50 mM Tris-Cl, pH 7.4, 50 mM NaF, containing 1% (v/v) Igepal CA-630 and supplemented with protease inhibitors). Cells were incubated for 30 minutes at 4°C with the lysis buffer and sonicated three times. Total cellular protein was measured using the BCA Protein Detection Kit (ThermoFisher, #23227) and 25 μg of cell lysate were electrophoresed at 100 V constant on NuPAGE 4– 12% Bis-Tris Midi gels (Invitrogen, NP0321BOX) with NuPage MES-SDS running buffer (Invitrogen, NP0002). After electrophoreses, gels were electroblotted in nitrocellulose membranes, using a wet-transfer approach. After electroblotting, the membranes were briefly rinsed in ultrapure water and incubated in 4% paraformaldehyde/PBS for 30 min at room temperature, followed by blocking for 1 hour at room temperature in TBS-Tween 0.1% (v/v) supplemented with 5% (w/v) dry milk. After blocking, membranes were incubated with primary antibodies against monomeric α-synuclein (BD Bioscience, #610787), multimeric α-synuclein (Sigma, ABN2265M), and β-actin (CellSignaling, #3700) or vinculin (Cell Signaling, #13901) at 4°C overnight. The next day, membranes were washed 3 × 10 min in PBS-Tween 0.1% and incubated with the corresponding secondary antibodies (anti-Mouse HRP, ThermoFisher, #31430 and anti-Rabbit HRP, ThermoFisher, #A24537) at RT for 1 h in the dark. Membranes were rinsed in PBS-Tween 0.1%, washed at least 3 × 10 min in PBS-Tween 0.1%, incubated for 3 minutes with ECL western blotting substrate, and imaged on a LI-COR Odyssey XF imaging system.

### Measurement of intracellular calcium levels

Intracellular calcium levels were evaluated using the cell-permeant calcium indicator Rhod-FF AM (Invitrogen, R23983). Briefly, 70×10^3^ mDA neurons were seeded per well in a Geltrex-coated black, clear-bottom 96-well cell culture plate and treated with the selected compound for three days. On neuronal maturation day 33, the neuronal medium was removed, and cells were washed with the pre-warmed HBS buffer (145 mM NaCl, 5 mM KCl, 1 mM CaCl_2_, 1 mM MgCl_2_, 10 mM HEPES, 10mM glucose) and incubated for 1 hour at room temperature, protected from light, with HBS buffer containing 0.1% (w/v) BSA and 5 μΜ Rhos-FF AM. After incubation, the cells were washed twice with 100μL per well HBS and were incubated for 20 minutes at room temperature, protected from light, for dye de-esterification. Fluorescence was measured 4 times, every 30 seconds, using a Cytation 5 (Bio-Tek) fluorescence plate reader, with excitation at 550 nm and emission at 580 nm, at 28°C. After the first measurement, HBS buffer supplemented with 10 μΜ of Ionomycin (Sigma, #19657) was added to the cells, and the plate was measured again (8 times (8 times, every 30 seconds). Fluorescence intensity was normalized to total cell protein levels, using the BCA Protein Detection Kit (ThermoFisher, #23227). Statistical analyses were conducted on data from three independent biological replicates.

### Evaluation of proteasomal activity

The proteasomal activity was evaluated by measuring the chymotrypsin-like and caspase-like activity. The chymotrypsin-like activity was measured using the fluorogenic substrate Suc-Leu-Leu-Val-Tyr-AMC (Enzo, BML-PB02-0005) and the caspase-like activity was measured by using the fluorogenic substrate Z-Leu-Leu-Glu-AMC (Enzo, BML-ZW9345-0005). Briefly, 60×10^3^ mDA neurons were seeded per well in a Geltrex-coated black, clear-bottom 96-well cell culture plate and treated with the selected compound for three days. On neuronal maturation day 33, the neuronal medium was removed and cells were washed with the pre-warmed assay buffer (20 mM Tris-HCl pH=7.5, 5 mM MgCl_2_, 0.1 mM EDTA, 1 mM DTT, 1 mM ATP, pH = 7.4). After washing, cells are incubated for 30 minutes at 37°C with 100 μL per well of assay buffer containing 100 μΜ of the respective fluorescent substrate. Finally, fluorescence was measured using a Cytation 5 (Bio-Tek) fluorescence plate reader, with excitation at 380 nm and emission at 460 nm, at 37°C. Fluorescence intensity was normalized to total cell protein levels, using the BCA Protein Detection Kit (ThermoFisher, #23227). Statistical analyses were conducted on data from three independent biological replicates.

### Structural analogues

Compounds were directly obtained from Selleckchem (CIDs: 5614, 2044, 1838043, 135419190), Enamine (CID: 135407924), or custom synthesized by SynHet (CID: 3511020). For compound CID 3511020, we chose the abbreviation CEPPE based on its IUPAC name 2-cyano-3-(3-ethoxy-4-propan-2-yloxyphenyl)prop-2-enethioamide. Chemical structures were plotted from their Simplified molecular-input line-entry system (SMILES) strings using Python. The rdkit package was used to create molecule objects from the SMILES strings and to compute 2D coordinates for these molecules. Tanimoto coefficients were calculated using their CID code via the PubChem Score Matrix service available at https://pubchem.ncbi.nlm.nih.gov/score_matrix/score_matrix.cgi.

## Acknowledgements

This research was funded by grants from the Fond National de Recherche (FNR) within the BRIDGES program “MOTASYN” (No.12719684). Fraunhofer ITMP received funding from European Joint Program on Rare Diseases (German Federal Ministry of Education and Research (BMBF), 01GM2002B). Ksilink was supported by the Programme d’investissements d’avenir (PIA) by the Secrétariat général pour l’investissement (SGPI) of the French government. We thank Dr. Tilo Kunath (University of Edinburgh) for providing the AST23 and AST23-2KO iPSC lines. We are grateful for the constant discussions with various colleagues at Ksilink and the Luxembourg Centre for Systems Biomedicine (LCSB) which improved this work.

## Author contributions

V.G. designed biochemical experiments, performed neuronal differentiation, biochemical assays, and data analysis; A.We. designed imaging experiments, performed neuronal culturing, imaging assay automation, and compound screening; L.C. developed and performed image analysis and data processing; D.H. performed MEA experiments; K.S. analyzed the MEA data; A.O. co-supervised data processing, P.A.B. co-initiated the study and contributed to grant writing; B.F.R.S. contributed to biochemical experiments; I.B. co-designed biochemical experiments; A.Wi., A.Z. and O.P. identified and provided Tyrphostin A9 structural analogues; R.K. co-initiated the study and supervised biochemical experiments; P.S. co-initiated the study and supported assay design; J.H.W. supervised the study, designed experiments, conceptualized and performed image analysis and data processing, and wrote the manuscript with input from all authors.

## Data availability statement

The datasets supporting the conclusions of this article are available in the GitHub repository, https://github.com/Ksilink/Notebooks/tree/main/Neuro/PD_MorphProfileScreening. Raw data is available as CSV files which can be processed using the provided Python-based Jupyter Notebooks.

## Supplementary Information

**Figure S1 (referring to Figure 1):**
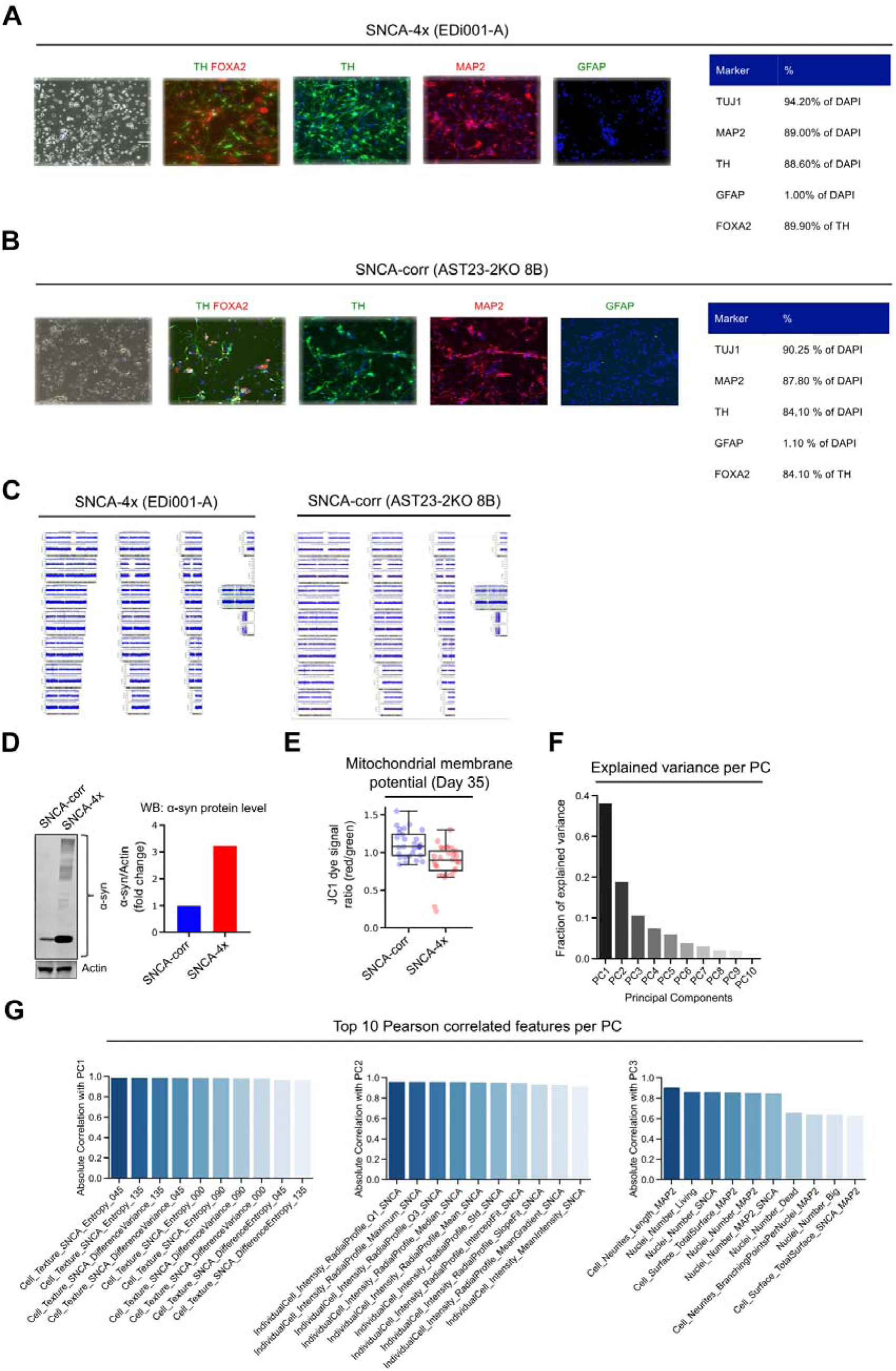
Quality control of neuronal cell lines for morphological profiling. **(A-B)** Immunofluorescent quantification of mDA SNCA-4x SNCA-corr neuron markers after differentiation D30. **(C)** Karyotype comparison between source iPSCs and D30 neurons. **(D)** Western blot densiometric quantification of αSyn signal from D70 mDA neuron lysates. The experiment was performed once. **(E)** Mitochondrial membrane potential assessment by JC1 dye fluorescent signal change in D35 neurons. Each data point shows technical replicates from a single experiment. The boxplots show the median and the 2nd and 3rd quartiles of the data. Whiskers indicate 1.5X of the interquartile range. **(F)** Explained data variation by first ten principal components (PCs) derived from principal components analysis. Pooled data from four independent experiments each containing at least 12 technical replicates per condition was used for analysis. **(G)** Top ten Pearson correlated morphological features for each of the first three PCs after PCA. Pooled data from four independent experiments each containing at least 12 technical replicates per condition was used for analysis.

**Figure S2 (referring to Figure 2):**
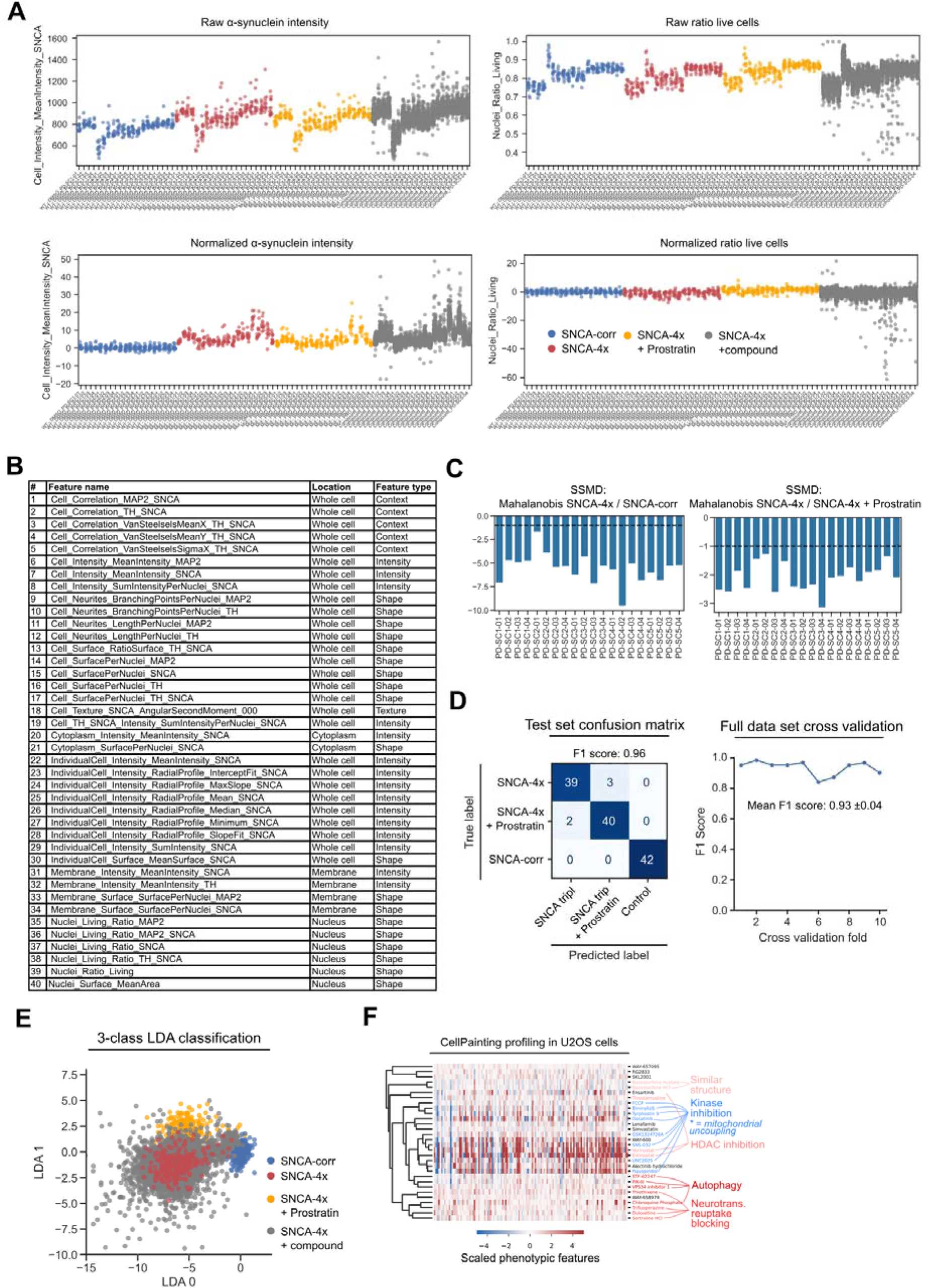
Primary screen quality assessment. **(A)** Raw and normalized αSyn staining intensity and live cell ratio values for all 20 screened plates. Each data point corresponds to the median value per well. **(B)** Selected features for hit identification based on <0.95 Pearson correlation. **(C)** Strictly standardized mean difference (SSMD) values per plate. The SSMD was calculated based on the morphological profile Mahalanobis distance between SNCA-4x + DMSO and SNCA-corr + DMSO conditions per plate (left panel) and between SNCA-4x + DMSO and SNCA-4x + Prostratin conditions per plate (right panel). SSMD values <-1 indicate a good plate quality (indicated by the dashed line). **(D)** Confusion matrix describing the per-class performance of the LDA classifier on the test data set containing 20% of the overall data not seen during training (left panel). Cross validation plot showing the LDA classifiers performance across 10 different slices of the full dataset (right panel). **(E)** Supervised Linear Discriminant Analysis (LDA) classification of Control neurons, and SNCA-4x neurons treated with either DMSO, Prostratin or screened compound. Each datapoint represents data from one well and for compounds the mean value of all four replicates is shown. **(F)** CellPainting profiles obtained in U2OS cells of hits selected during primary screening clustered by their Cosine similarity. Compounds are color coded by attributed biological activity.

**Figure S3 (referring to Figure 3):**
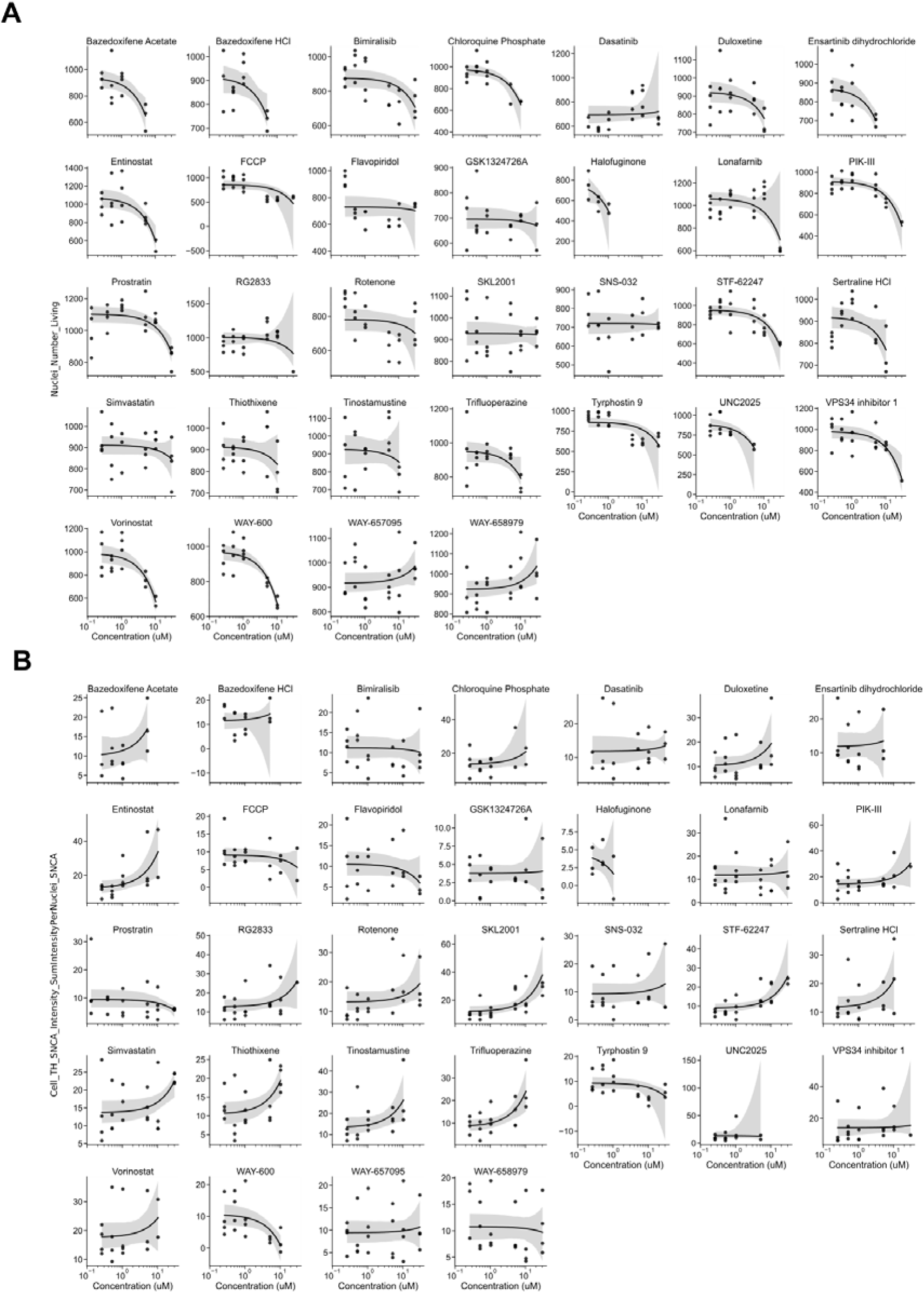
Dose-response assessment of cell viability and αSyn level. **(A)** Cell viability assessment based on Hoechst-stained nuclear size and intensity. mDA neurons were treated with 0.25-20 μM compound for 72 hours before fixation. **(B)** Immunostaining-based αSyn level assessment. Toxic compound concentrations were excluded from the graph. Each compound concentration was tested at least in triplicate. The mean and 95% confidence interval is shown.

**Figure S4 (referring to Figure 3):**
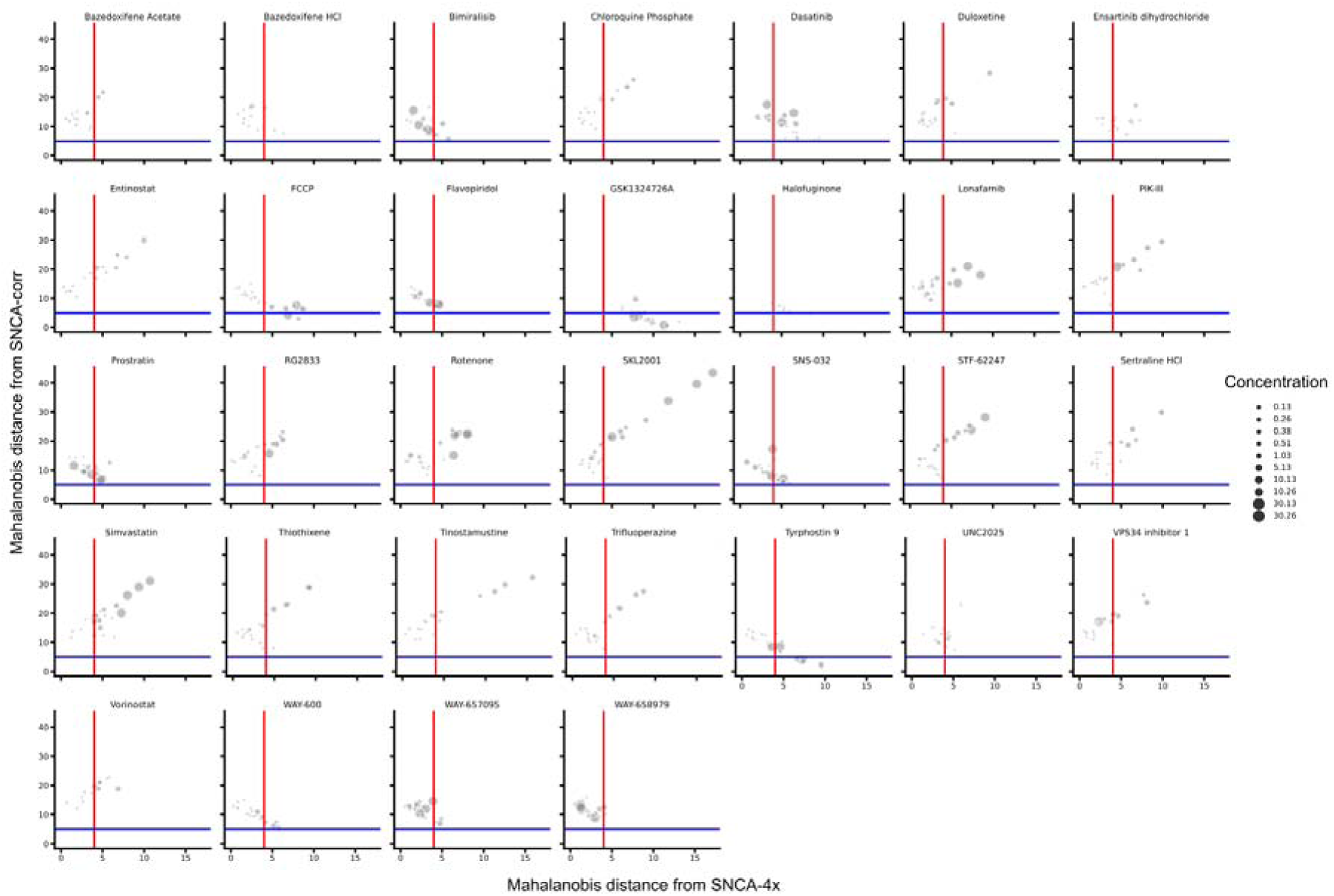
Dose-response assessment of compound effects on morphological profiles. Morphological profile Mahalanobis distances of compound treated mDA neurons relative to SNCA-4x and SNCA-corr morphological profiles. The redline line indicates the 4σ window from the SNCA-4x median and blue line the indicates the 2.5σ window from the SNCA-corr median. Dot sizes represent the used concentration. Each compound concentration was tested at least in triplicate. Mean and 95% confidence interval are shown.

**Figure S5 (referring to Figure 4):**
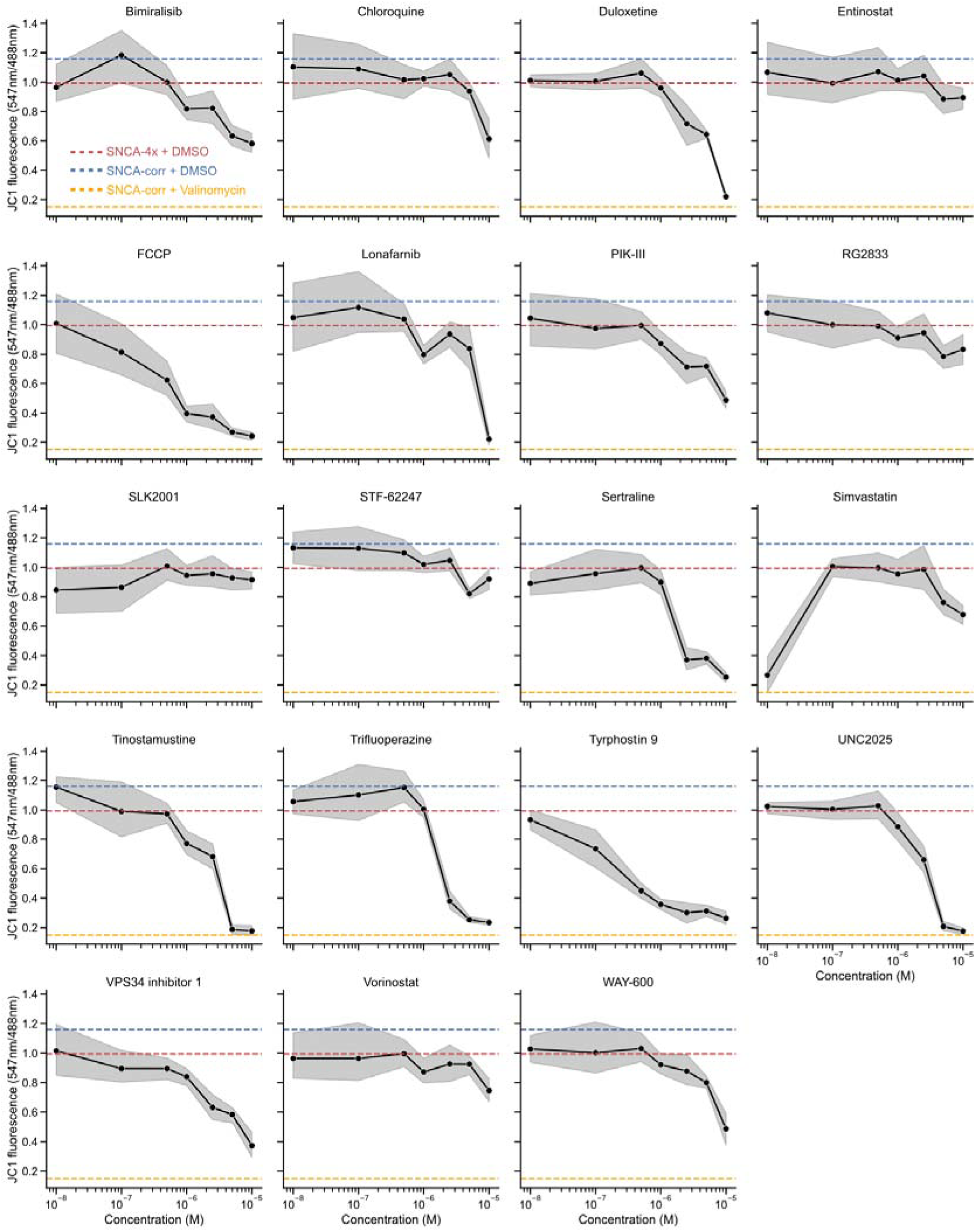
Dose-response assessment of compound effects on mitochondrial membrane potential. mDA neurons were treated with 0.1-10 μM of each compound for 72 hours and the mitochondrial membrane potential was measured using the JC1 dye. Three independent experiments with at least three technical replicates each were performed. The mean and 95% confidence interval is shown. Dashed lines indicate the means of the indicated control treatments.

**Figure S6 (referring to Figure 4):**
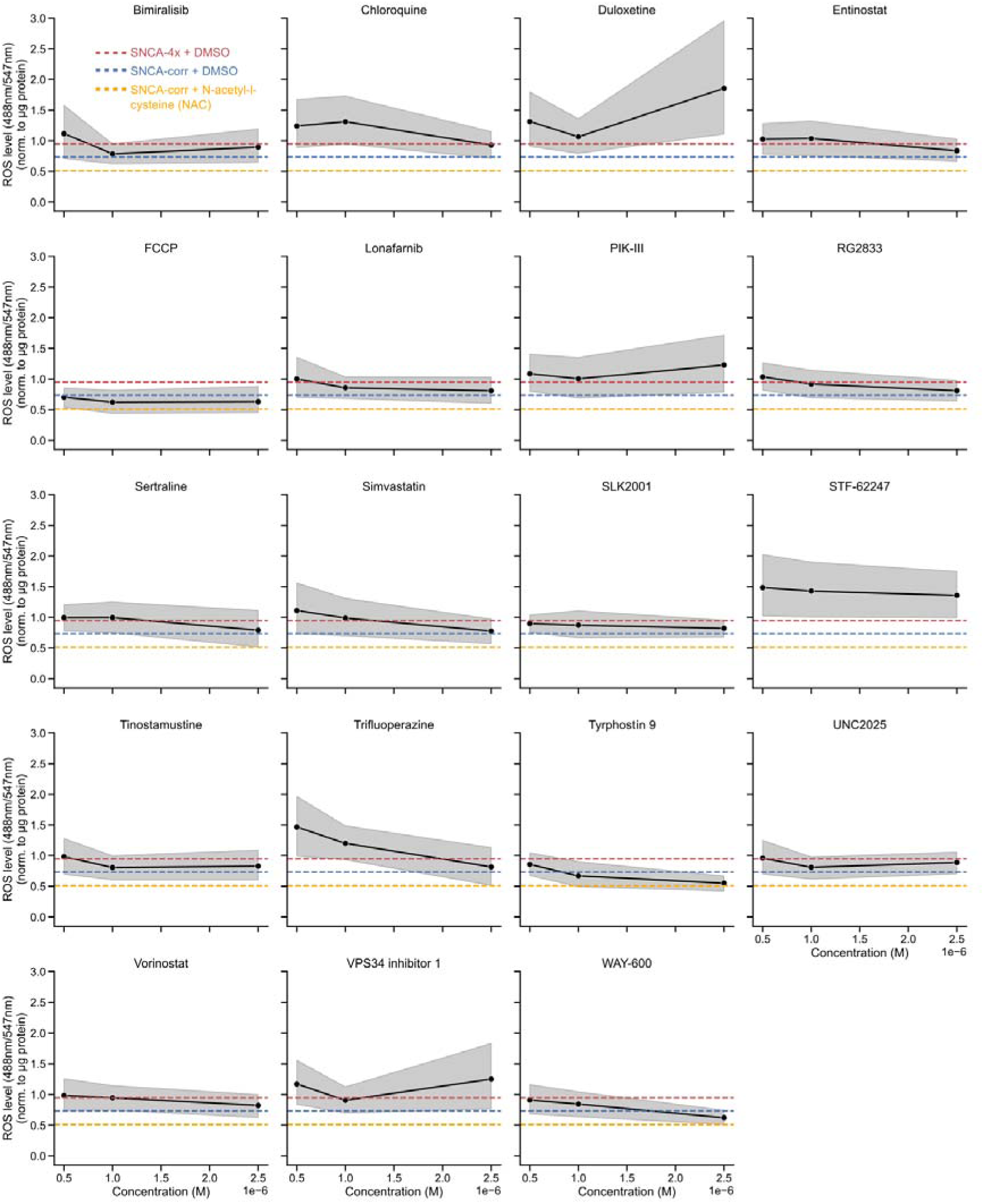
Dose-response assessment of compound effects neuronal ROS level. mDA neurons were treated with 0.5, 1, and 2.5 μM of each compound for 72 hours and the ROS level was determined using the ROS-ID Total ROS/Superoxide detection kit. The obtained signal was normalized to total protein content. Three independent experiments with each at least three technical replicates were performed. The mean and 95% confidence interval is shown. Dashed lines indicate the means of the indicated control treatments.

**Figure S7 (referring to Figure 5):**
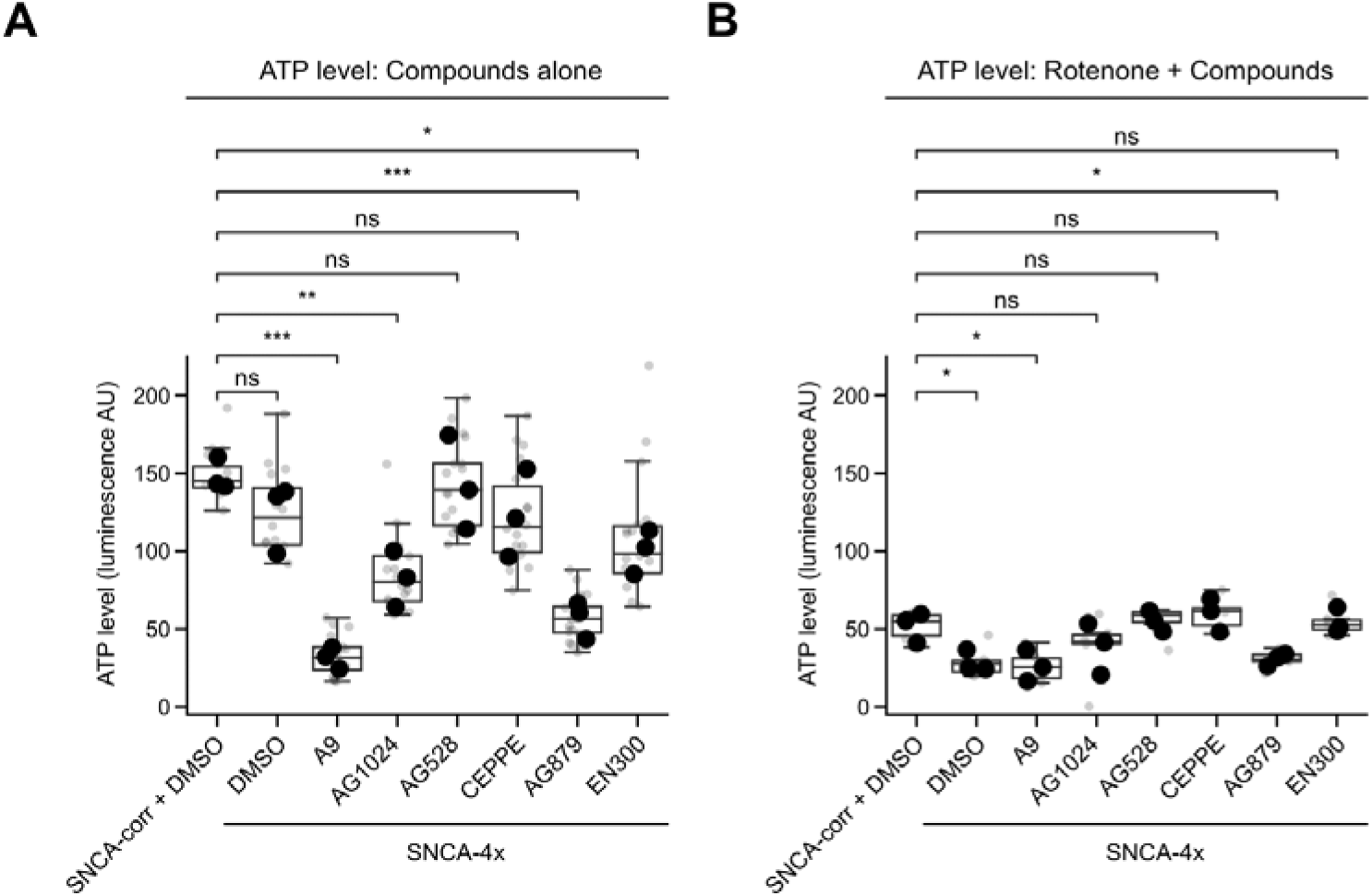
Tyrphostin A9 structural analogues with mitochondrial uncoupling activity reduce cellular ATP levels. **(A)** SNCA-4x and SNCA-corr mDA neurons were differentiated until D33, treated with DMSO, 2.5 μM Tyrphostin A9 or its structural analogues for 72 hours and ATP levels were determined in a luminescence-based plate reader assay. **(B)** Same as in (A), but mDA neurons were simultaneously incubated with compound and the mitochondrial complex I inhibitor Rotenone. Three independent differentiations with at least three technical replicates each were performed. Large markers indicate biological replicate medians and small markers represent all technical replicate data. The boxplots show the median and the 2nd and 3rd quartiles of the data. Whiskers indicate 1.5X of the interquartile range. Welch’s unequal variances t-test was performed using the medians of all biological replicates. ns = not significant, * = p < 0.05, ** = p < 0.01, *** = p < 0.001.

**Figure S8 (referring to Figure 6):**
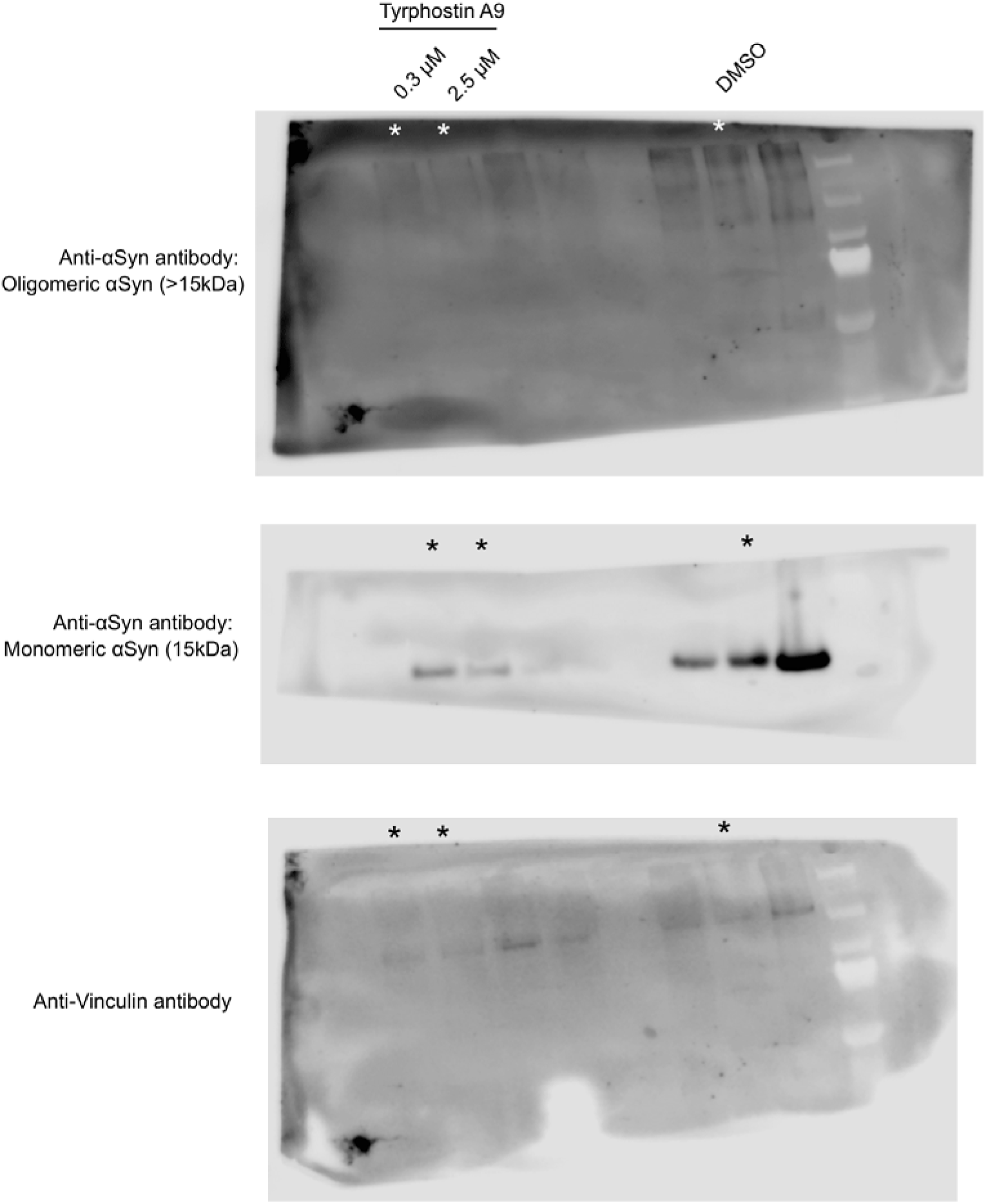
Full gel/blot images. Shown are raw uncropped images of membranes stained with antibodies against αSyn and Vinculin.

**Table S1:** Supplementary dataset of identified compounds after morphological profiling including morphological features.

**Table S2:** Key resource table.

**Table S3:** Supplementary dataset: Compound library overview.

**Table S4:** Supplementary dataset: Morphological feature overview.

